# Structural basis for the mechanism and antagonism of receptor signaling mediated by Interleukin-9 (IL-9)

**DOI:** 10.1101/2022.12.30.522308

**Authors:** Tatjana De Vos, Marie Godar, Fabian Bick, Anna C. Papageorgiou, Thomas Evangelidis, Iva Marković, Eva Mortier, Laure Dumoutier, Konstantinos Tripsianes, Christophe Blanchetot, Savvas N. Savvides

## Abstract

Interleukin-9 (IL-9) is the hallmark cytokine in Th9 immunity and is also central to Innate Lymphocyte 2 (ILC2) biology. Furthermore, receptor signaling mediated by IL-9 has been linked to inflammatory and autoimmune diseases, and cancer. Despite its functional pleiotropy, the structure-function landscape of IL-9 had remained enigmatic. Here, we show via a combination of X-ray crystallography and NMR that human IL-9 adopts a helical bundle fold with unprecedented structural features among helical cytokines, including five disulfide bridges. Binding of IL-9 to the interdomain junction of IL-9Rα results in marked structural changes on the opposite face of IL-9 that prime the binary complex for recruiting the common gamma chain (γc) for signaling. Surprisingly, this tripartite cytokine-receptor assembly displays a markedly lower affinity than the IL-9: IL-9Rα complex, which we trace to distinct features of IL-9Rα that might destabilize the ternary complex. Furthermore, we developed monoclonal antibodies that antagonize IL-9 activity by sterically competing for the binding footprint of IL-9Rα. Collectively, we here provide a structural and mechanistic blueprint to facilitate interrogation and modulation of pleiotropic signaling outputs of IL-9 in physiology and disease.

## Introduction

Originally, IL-9 was discovered in 1988 as a T-cell growth factor labeled P40 due to its apparent molecular weight (*1*), and in 1989 as a factor with mast-cell growth enhancing activity (MEA) (*2, 3*). Because of its pleiotropic effects on myeloid and lymphoid cells, it was eventually renamed interleukin-9 (*4*). Interleukin-9 (IL-9) was found to signal by binding to a heterodimeric receptor consisting of the specific IL-9 receptor α (IL-9Rα) and the common gamma chain (γc) which is shared with IL-2, IL-4, IL-7, IL-15 and IL-21. The binding of IL-9 to its receptors, activates the associated Janus Kinases (JAK)1 and JAK3, resulting in the activation of signal transducer and activator of transcription (STAT)1, STAT3 or STAT5 pathways (*5*), the mitogen-activated protein kinase (MAPK) pathway (*6*), and insulin-related substrate (IRS) pathway (*5, 7*). However, it was long disregarded as just another T helper 2 (Th2)-type cytokine. This changed when a new distinct subset of T helper (Th) cells was identified with IL-9 as their signature cytokine, referred to as Th9 cells (*8, 9*). In the wake of that, more IL-9-producing cells were discovered or firmly acknowledged, including innate lymphoid cells 2 (ILC2) (*10*), Th17 cells (*11*), mast cells (*12, 13*), osteoblasts (*14*), NKT cells (*15*) and memory B cells (*16*).

This wide range of cellular sources points to a complex system of IL-9 expression and suggests the involvement of IL-9 in multiple physiological conditions and diseases. IL-9 has a long history as an aggravating agent in asthma (*17*–*20*) in which it is associated with mucus production, bronchial hyperresponsiveness and pulmonary mastocytosis (*19*). Recently, IL-9 was identified as the most upregulated gene in HDM-reactive Th2 cells in allergic asthmatic patients in a single-cell transcriptomics study (*21*). Furthermore, it has negative effects in multiple hematological malignancies including anaplastic large cell lymphoma (*22*) and Hodgkin’s disease (*23*), where the high expression of IL-9Rα on the cancer cells results in the promotion of their survival and proliferation. On the other hand, IL-9 also has established anti-tumor properties in melanoma (*24, 25*). Adoptive transfer of Th9 cells or addition of recombinant IL-9 resulted in reduced melanoma growth and increased survival (*24*). Next to this, IL-9 has been shown to be a major factor in the resolution of inflammation in a chronic model for rheumatoid arthritis, where IL-9-primed ILC2s activate regulatory T (Treg) cells (*26*). It has also been suggested as a promising candidate for chemotherapy-induced thrombocytopenia (CIT) treatment since IL-9 produced by osteoblasts in the bone marrow supports megakaryopoiesis resulting in higher levels of platelets and enhanced recovery in vivo (*14*).

This wave of biomedical findings has been met with a paucity in structural and mechanistic information. Efforts have been made to model the cytokine in order to shine a light on this signaling complex (*27, 28*). Here, we present the structure of human IL-9 (hIL-9) in solution and of hIL-9 bound to the extracellular domain of its specific hIL-9Rα receptor. We show that IL-9 binding to hIL-9Rα occurs with high affinity and leads to a drastic restructuring of hIL-9. The recruitment of hγc to the binary hIL-9:hIL-9Rα complex occurs with severely reduced affinity, which we trace to distinctive features in hIL-9Rα that might compromise the binding of hγc. Additionally, we present several anti-hIL-9 antibodies which antagonize IL-9 activity by competing against hIL-9Rα for binding to IL-9. Together our findings provide the missing structural framework for interrogating the pleiotropic signaling outputs of IL-9 in physiology and disease.

## Results

### Human IL-9 binds to IL-9Rα with high affinity

To study the thermodynamic and kinetic profile underlying the assembly of the IL-9 signaling complex we used a range of different constructs covering the recombinant production of mature hIL-9 (residues 19-144) in both *E. coli* and HEK293T cells, the extracellular domain of hIL-9Rα (residues 40-261) in *E. coli*, the trimmed extracellular domain of hγ_c_ (residues 56-254) in Sf9 cells, and the full extracellular domain of hγ_c_ (residues 23-262) in HEK293T cells (**Fig. S1**). First, we confirmed the proper folding of recombinant hIL-9 produced in HEK293T cells and in *E. coli*, following its in vitro refolding from inclusion bodies based on an in-house protocol (*29*), by testing the activity in a cell proliferation assay using the human megakaryocytic leukemic cell line MO7e that is responsive to IL-9. Our results confirm that both recombinantly produced proteins are active and therefore appropriate to use for subsequent studies (**Fig. 1a**). Next, we focussed on quantifying the kinetics, thermodynamics, and affinity of the binary hIL-9:hIL-9Ra complex using biolayer interferometry (BLI) and isothermal titration calorimetry (ITC). This interaction is entropically-driven binding mode with low nanomolar affinity (*K*_D_ = 20 ± 3 nM via BLI, *K*_D_ = 9 ± 9 nM via ITC) (**Fig. 1b,c**). The low nanomolar affinity is achieved through a fast association rate constant (*k*_a_ = 5 ± 2 × 10^5^ M^-1^s^-1^) and a fast dissociation rate constant (*k*_d_ = 1.1 ± 0.4 × 10^-2^ s^-1^). To enable better comparisons of affinities obtained by ITC and BLI we repeated the ITC measurements at 298 K to match the temperature of the BLI measurements (**Fig. S2a**). Interstingly, the change in temperature causes a change in the apparent enthalpy/entropy ratio resulting in endothermic binding events, but still leads to a high affinity interaction of the binary hIL-9:hIL-9Rα complex (*K*_D_ = 4 ± 4 nM). Thus, the binary complex comprising human IL-9 and IL-9Rα assembles with high affinity and an interaction stoichiometry of 1:1.

**Figure 1:**
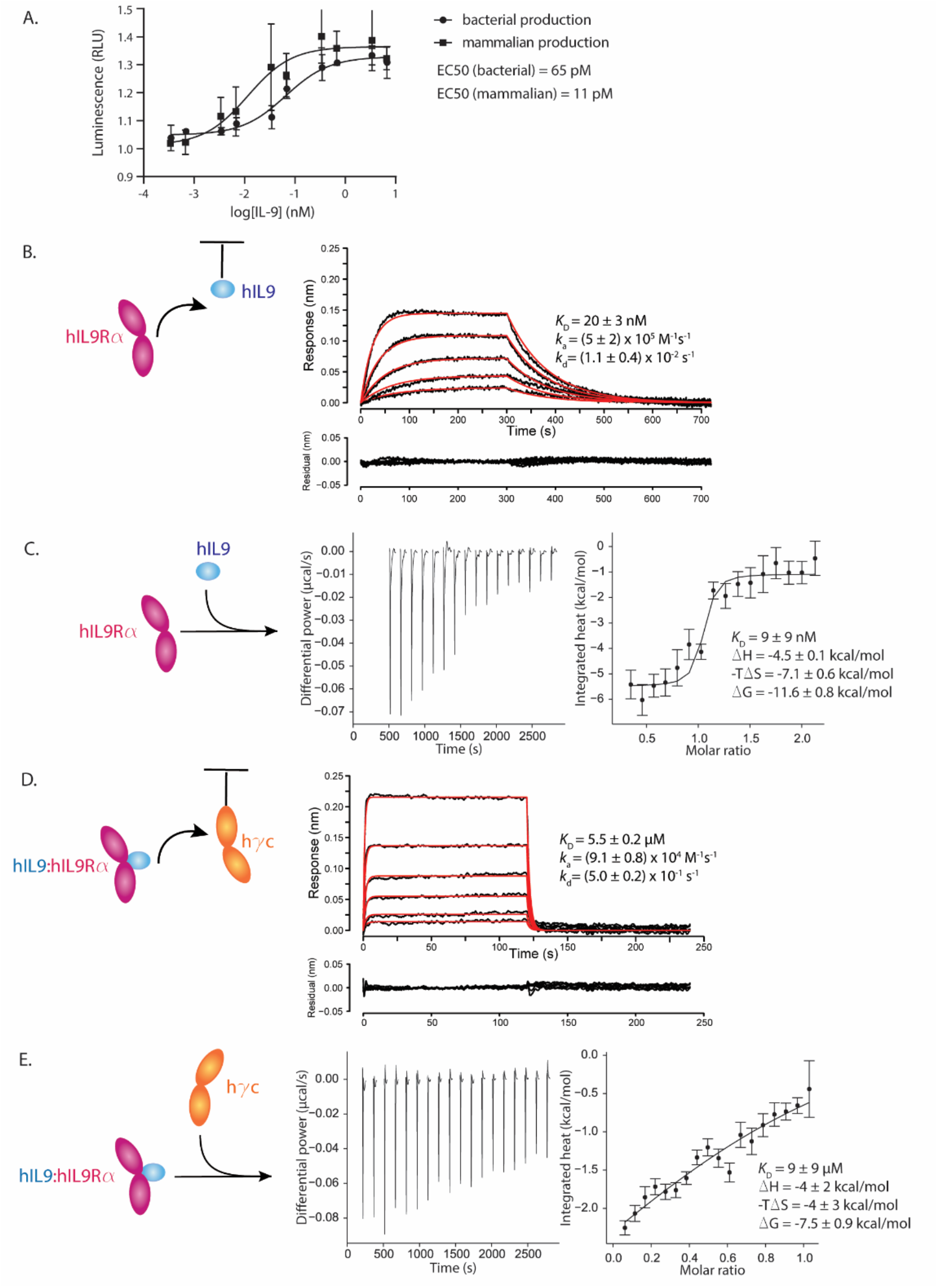
The human IL-9Rα:IL-9:γc ternary complex shows a decline in affinity compared to the IL-9Rα:IL-9 binary complex. (A) Proliferation assay of MO7e cells upon stimulation with increasing concentrations of glycosylated hIL-9 produced in HEK293T cells or non-glycosylated hIL-9 produced in *E. coli*. Assays were carried out in triplicates and the represented values are averaged values from three independent experiments and their s.d. Representative experiment is shown. (B,C,D,E) Kinetic and thermodynamic characterization of the binary hIL-9:hIL-9Rα complex and the ternary hIL-9:hIL-9Rα:hγc complex. (B) Representative BLI data traces fitted with a 1:1 binding model to quantify the kinetics and binding affinity of hIL-9Rα (highest concentration of 49 nM, two-fold dilution series) to coupled hIL-9 from HEK293T cells. The represented values are averaged values and their s.d. from four replicate experiments. (C) Representative ITC experiment of the titration of 58 μM non-glycosylated hIL-9 into 5.1 μM hIL-9Rα with a local correction factor of 0.97. The represented values are averaged values and their s.d. from two replicate experiments. (D) Representative BLI data traces fitted with a 1:1 binding model to quantify the kinetics and binding affinity of hIL-9:hIL-9Rα containing glycosylated hIL-9 (highest concentration of 6,5 μM, two-fold dilution series) to coupled hγc. The represented values are averaged values and their s.d. from three replicate experiments. (E) Representative ITC experiment of the titration of 172 μM hγ_c_ into 22.5 μM non-glycosylated hIL-9:hIL-9Rα with a local correction factor of 0.64. The represented values are averaged values and their s.d. from three replicate experiments.

### Recruitment of the γ_c_ receptor weakens the affinity of the receptor complex mediated by IL-9

We were surprised to measure a The hIL-9:hIL-9Rα interaction is in stark contrast with the binding mode of the ternary hIL-9:hIL-9Rα:hγc complex where the affinity is dramatically lowered to the low micromolar range (*K*_D_ = 5.5 ± 0.2 μM by BLI, *K*_D_ = 9 ± 9 μM by ITC) with a poor association rate (*k*_a_ = 9.1 ± 0.8) × 10^4^ M^-1^s^-1^) and dissociation rate (*k*_d_ = 5.0 ± 0.2 × 10^-1^ s^-1^) (**Fig. 1d,e**). This mechanistically intriguing observation applies to the interaction between hγ_c_ and the binary hIL-9:hIL-9Rα complex applies to both the use of glycosylated or non-glycosylated hIL-9 (**Fig. S2b**). We also performed the same experiments with the complete ectodomain of hγc (residues 23-262), to investigate whether the trimmed N-terminus of hγc is involved in this interaction (**Fig. S2c,d**). These experiments show no difference from the experiments with the trimmed hγc (residues 56-254), indicating that the N-terminal 32 amino acids are not involved in the binding of the binary hIL-9:hIL-9Rα complex. As discussed above, all ITC experiments were repeated at 298 K which confirmed the main insights (**Fig. S2a,e**). The binary hIL-9:hIL-9Rα complex binds with low nanomolar affinity whereas the ternary hIL-9:hIL-9Rα:hγc complex drops to a *K*_D_ = 2.5 ± 0.4 μM. We also probed the other binary interactions between the single components by ITC. We investigated whether unbound hIL-9 has an affinity for hγ_c_, whether the two receptor chains have an affinity towards each other, and whether the hIL-9Rα chain might self-associate. These experiments were performed both at 310 K and at 298 K, but we were unable to detect binding for any of the interrogated binder combinations (**Fig. S3**). Since for other members of the IL-2 cytokine family weak binding between the specific receptor and γ_c_ has been demonstrated, we performed additional ITC measurements with the receptor components in TRIS buffer to amplify possible enthalpic signal (**Fig. S3i**). However, we were again unable to detect any measurable binding between hIL-9Rα and hγ_c_ in this experimental setup.

### Human IL-9 is a flexible helical cytokine with unprecedented features

We show that hIL-9 engages hIL-9Rα with a high affinity, but that the subsequent binding of hγ_c_, resulting in the assembly of the ternary complex, occurs with low affinity. By tackling the structural basis of the hIL-9 signaling complex, we aim to elucidate this sequence of events. To take an in-depth look at the unbound cytokine we initiated the structure determination of hIL-9 in solution by NMR (**Table 1**) (*30*). To perform these experiments, we expressed ^13^C, ^15^N double-labeled hIL-9 in *E. coli* in minimal media complemented with isotope-labeled compounds, followed by in vitro refolding of the inclusion bodies. Mature hIL-9 follows the paradigm of the IL-2 family where it adopts a four helical bundle fold with an up-up-down-down topology (**Fig. 2a**). The four helices are connected through two long overhead loops AB and CD, and a shorter BC-loop. A remarkable feature of this cytokine is the rather short B-helix, which additionally displays an unusually high degree of flexibility, together with the long AB- and CD-loop (**Fig. 2b**). This flexibility is remarkable given the tight locks provided by the ten cysteines that are all engaged in disulfide bridges (**Fig. 2a**). These cysteines are highly conserved among IL-9 orthologs (Fig. S4a). Another aspect of the flexibility displayed by this cytokine is reflected in the D-helix, which shows a slight sway of 5-10° across the 20 states of the NMR ensemble (**Fig. 2a**).

**Table 1.** NMR statistics for 20 models of human IL-9.

| NMR distance and dihedral restraints | hIL-9 |
| --- | --- |
| Distance restraints |  |
| Total NOE | 2295 |
| Intra-residue ($i=j$) | 327 |
| Sequential ($\vert i-j\vert = 1$) | 708 |
| Medium-range ($1 < \vert i-j\vert < 5$) | 828 |
| Long-range ($\vert i-j\vert \geq 5$) | 432 |
| Total dihedral angle restraints | 190 |
| $\phi$, $\psi$ | 92, 98 |
| Total disulfide bond restraints | 5 |
| Violations (mean ± s.d.) | |
| Distance restraints ($\text{\AA}$) | $0.019 \pm 0.001$ |
| Dihedral angle restraints ($^\circ$) | $1.552 \pm 0.064$ |
| Max. dihedral angle violation ($^\circ$) | 9.80 |
| Max. distance constraint violation ($\text{\AA}$) | 0.356 |
| Deviations from idealized geometry |  |
| Bond lengths ($\text{\AA}$), angles ($^\circ$) | $0.011 \pm 0.000$, $1.27 \pm 0.030$ |
| Improper angles ($^\circ$) | $1.48 \pm 0.074$ |
| Average pairwise r.m.s. deviation* ($\text{\AA}$) | |
| Backbone, Heavy | $0.60 \pm 0.12$, $0.97 \pm 0.13$ |
| Ramachandran plot statistics (%) |  |
| favored regions | 95.5 |
| allowed regions | 4.4 |
| generously allowed regions | 0.1 |
| disallowed regions | 0.0 |
| PDB | 7OX6 |
| BMRB | 34640 |

\* Ordered residues 19-44, 55-64, 69-140

**Figure 2:**
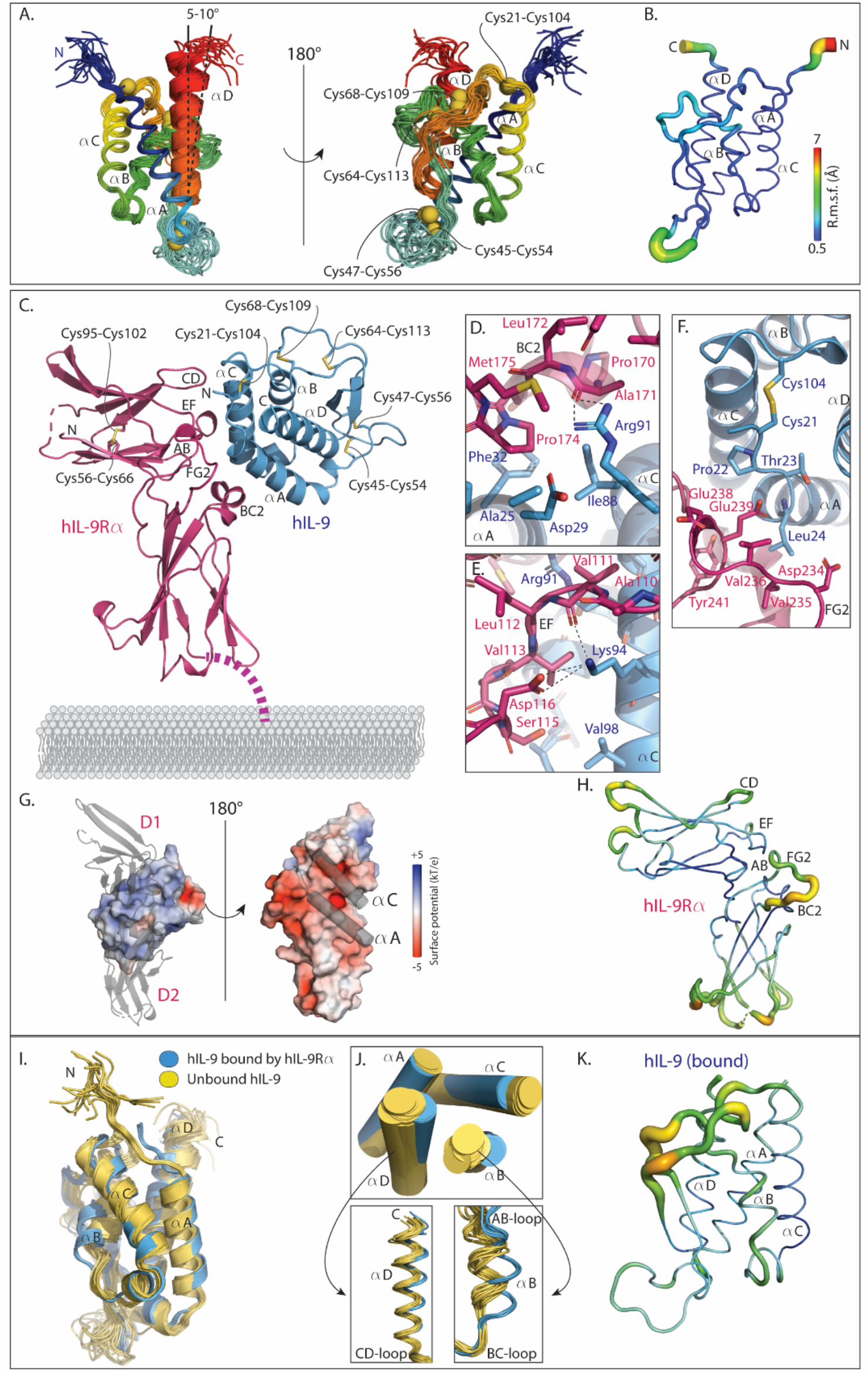
hIL-9 in solution *versus* hIL-9 bound to hIL-9Rα. (A) Cartoon representation of ensemble (20 states) of hIL-9 in solution as determined by NMR. Cysteine pairs are represented as yellow spheres from 1 state. Blue-red color gradient from N- to C-terminus. (B) Root mean square fluctuation (r.m.s.f.) analysis of hIL-9. The residues are colored according to their r.m.s.f values. The radius of the cartoon loop varies according to the r.m.s.f. value. (C) Cartoon representation of the determined crystal structure of the hIL-9:hIL-9Rα complex. (D) Close-up view of the hIL-9:hIL-9Rα interaction in the vicinity of Arg91 in hIL-9. (E) Close-up view of the hIL-9:hIL-9Rα interaction in the vicinity of Lys94 in hIL-9. (F) Close-up view of the interaction between the N-terminal part of hIL-9 and the FG2-loop of hIL-9Rα. (G) Complementary electrostatic surface potential of hIL-9 and hIL-9Rα. (H) Representation of hIL-9Rα (chain A) with colors and radius of the cartoon varying according to the B-factors. (I) Comparison between the ensemble of unbound hIL-9 as determined by NMR with the 8 copies in the ASU of bound hIL-9 as determined by X-ray crystallography. (J) Comparison of the helical bundles of unbound hIL-9 and bound hIL-9 with close-up views of the B- and D-helix. (K) Representation of hIL-9 (chain B) with colors and radius of the cartoon varying according to the B-factors.

### Structural basis of the human IL-9:IL-9Rα complex

To obtain structural insights into the assembly of the receptor complex mediated by human IL-9 we pursued structural studies by X-ray crystallography of human IL-9 in complex with hIL-9Rα and hIL-9Rα/hγ_c_, respectively. For this purpose we used recombinant hIL-9 (residues 19-144) and hIL-9Rα (residues 40-261) produced in *E. coli* (**Fig. S1a,b**) and the trimmed ectodomain of hγ_c_ (residues 56-254) produced in *Sf9* cells (Fig. S1d). We biochemically reconstituted and purified the stoichiometric hIL-9:hIL-9Rα binary and hIL-9:hIL-9Rα:hγc ternary complexes (Fig. S1c,e). As the ternary complex failed to produce well-diffracting crystals and due to the fact that N-glycan shaving resulted in a severe decrease in the solubility and stability of the complex, we shifted our efforts to the hIL-9:hIL-9Rα binary complex. Indeed, crystals of the hIL-9:hIL-9Rα complex proved of much better diffraction quality and enabled elucidation of the crystal structure of the hIL-9:hIL-9Rα complex to 3.1 Å resolution (**Table 2**).

**Table 2:** Crystallographic data and refinement statistics.

| Table 2: Crystallographic data and refinement statistics |  |  |  |  |  |
| --- | --- | --- | --- | --- | --- |
|  | hIL-9:hIL-9Ra | Fab 6E2:hIL-9 | Fab 7D6:hIL-9 | Fab 6D3:hIL-9 | Fab 35D8:mIL-9 |
| Crystallization conditions | 0.1M Tris, 0.1M ammonium sulfate, 18% PEG 1500, pH 5.6 | 0.2M lithium sulfate, 0.1M phosphate/citric acid, pH 3.8 | 0.25M ammonium nitrate, 18% PEG 3350, 5% ethylene glycol | 0.1M ammonium sulfate, 0.1M Tris, 16% PEG 1500, pH 5.8 | 0.2M Zinc acetate, 23% PEG 3350 |
| Protein concentration cryoprotectant | 8.5 mg/ml<br>20% ethylene glycol | 13.4 mg/ml<br>20% glycerol | 15 mg/ml<br>20% ethylene glycol | 14 mg/ml<br>25% PEG 1500 | 13.5 mg/ml<br>20% DMSO/ethylene glycol/glycerol (2:2:1) |
| Data Collection<sup>a</sup> |  |  |  |  |  |
| X-ray source | SOLEIL (Proxima 2A) | ESRF (ID23-1) | ESRF (ID30A-3) | PETRA III (P14) | PETRA III (P13) |
| Wavelength (Å) | 0.98 | 0.97242 | 0.96770 | 0.9763 | 0.9762 |
| Space group | P2<sub>1</sub> | P1 | P2<sub>1</sub> | P2<sub>1</sub>2<sub>1</sub>2 | C222<sub>1</sub> |
| Cell dimensions<br>a, b, c (Å) | 85.45, 107.81, 159.55 | 69.37, 94.37, 107.34 | 81.35, 102.75, 164.37 | 60.48, 81.87, 116.55 | 70.21, 78.92, 192.10 |
| α, β, γ (°) | 90, 90.76, 90 | 82.53, 72.00, 80.79 | 90, 94.12, 90 | 90, 90, 90 | 90, 90, 90 |
| Resolution (Å) | 50 – 3.1<br>(3.28 – 3.1) | 50- 3.35<br>(3.55- 3.35) | 50 – 2.5<br>(2.65 – 2.5) | 999-1.7<br>(1.79 – 1.7) | 999 – 1.8<br>(1.9 – 1.8) |
| R<sub>meas</sub> (%) | 9.7 (168) | 27.8 (112.1) | 11.4 (114.0) | 7.5 (162.6) | 13.1 (152.9) |
| <I/σ> | 11.39 (0.82) | 4.78 (1.04) | 11.99 (1.23) | 21.59 (1.66) | 9.49 (1.06) |
| CC<sup>1/2</sup> (%) | 99.8 (28.2) | 96.0 (51.7) | 99.7 (49.5) | 99.9 (70.3) | 99.7 (44.3) |
| Completeness (%) | 98.9 (94.8) | 93.6 (89.60) | 97.7 (95.9) | 99.2 (97.9) | 99.7 (99.1) |
| Redundancy | 3.5 (3.3) | 2.9 (2.8) | 3.2 (3.2) | 13.4 (13.6) | 5.8 (5.8) |
| Wilson B (Å<sup>2</sup>) | 100.794 | 56.48 | 55.16 | 36.21 | 35.04 |
| Refinement<sup>b</sup> |  |  |  |  |  |
| Resolution (Å) | 45.59- 3.09 | 48.92- 3.34 | 40.57- 2.49 | 41.96- 1.70 | 46.04- 1.80 |
| No. reflections (R<sub>free</sub>) | 52738 (2640) | 34587 (1728) | 92479 (4624) | 63802 (3188) | 49646 (2482) |
| R<sub>work</sub>/R<sub>free</sub> (%) | 21.75/27.61 | 22.58/27.83 | 21.00/26.19 | 17.61/21.02 | 18.33/22.87 |
| Non-Hydrogen atoms |  |  |  |  |  |
| Protein | 19758 | 16132 | 16289 | 4117 | 4087 |
| Ligand/ion | 50 | 25 | 0 | 25 | 23 |
| Water | 6 | 2 | 225 | 424 | 316 |
| R.m.s. deviations |  |  |  |  |  |
| Bond lengths (Å), angles (°) | 0.003, 0.73 | 0.005, 1.05 | 0.011, 1.36 | 0.010, 1.16 | 0.008, 0.97 |
| Ramachandran |  |  |  |  |  |
| favored | 94.12 | 95.31 | 94.61 | 98.32 | 97.94 |
| allowed | 5.39 | 4.31 | 4.82 | 1.68 | 1.68 |
| outliers | 0.50 | 0.38 | 0.57 | 0.00 | 0.37 |
| Rotamer outliers (°) | 0.09 | 0.00 | 0.00 | 0.00 | 0.00 |
| Clash score | 7.39 | 9.52 | 11.87 | 2.45 | 6.92 |
| B-factors (Å<sup>2</sup>) |  |  |  |  |  |
| Protein | 124.11 | 78.46 | 71.06 | 39.57 | 40.39 |
| Ligand/ion | 141.07 | 96.97 | 0 | 71.13 | 58.75 |
| Water | 100.24 | 25.75 | 52.14 | 45.12 | 42.41 |
| PDB accession codes | 7OX5 | 7OX2 | 7OX1 | 7OX3 | 7OX4 |

<sup>a</sup> Values reported by XDS. <sup>b</sup> Values reported by Phenix.

hIL-9 engages hIL-9Rα at the elbow tip of the cytokine-binding homology region (CHR) consisting of the two domains of the receptor, which are positioned at a 90° angle of each other (**Fig. 2c**). This interaction, with an average interface area of 849 Å^2^, is mainly driven by hydrophobic contacts (**Table S1**). This hydrophobic interface of the binary complex corresponds well with the high entropic contribution to the thermodynamic footprint determined by ITC. Some specificity by polar interactions is governed by the C-helix of hIL-9, more precisely by residues Arg91 and Lys94, interacting with residues on the BC2-loop and EF-loop of the receptor respectively (**Fig. 2d,e**). Additionally, the N-terminal end of hIL-9 is engaged by the FG2-loop of the receptor (**Fig. 2f**). Here, Glu239 of the FG2-loop might serve to stabilize the local dipole of the A-helix. In contrast to their hydrophobic interface, hIL-9 and hIL-9Rα display distinct complementary electrostatics on their binding interfaces, with the site on hIL-9 engaging hIL-9Rα being positively charged and the corresponding site on hIL-9Rα being largely negatively charged (**Fig. 2g**). These long-range interactions might be responsible for drawing the two proteins together and might contribute to the measured high affinity for this complex. To gain insights into the remaining flexibility of the receptor we examined the B-factors. Even though these vary across the 8 copies in the crystal asymmetric unit, the temperature factors within the molecules display similar fluctuations. This shows that the bound receptor retains flexibility near the outsides of the two domains (**Fig. 2h**). The binding sites in the AB-, EF-, and BC2-loop are clearly more restrained in flexibility (**Fig. 2h**) while in comparison, even though the CD- and FG2-loop are part of the interface, they still have high B-factors indicating an intrinsic flexibility of these patches (**Fig. 2h**).

### Binding of hIL-9Rα restructures hIL-9 to prime γc recruitment

To obtain insights into the possible conformational plasticity of IL-9 upon receptor binding we superposed the ensemble of unbound hIL-9 determined by NMR with the 8 copies of hIL-9 determined in the crystal structure of hIL-9 in its bound state to IL-9Rα. The AC-face of the cytokine, which carries the IL-9Rα binding footprint undergoes only limited restructuring at the end of helix C upon binding (**Fig. 2i,j**). This might be due to the direct binding of hIL-9Rα to this site or to the interaction of the receptor with the tip of the A-helix, which is linked to the C-helix through a disulfide bridge. Nevertheless, the AC-face of IL-9 remains rather unchanged between the bound and the unbound structures, suggesting that the binding site for hIL-9Rα is pre-formed. However, the largest structural shifts are visible at helices B and D on the opposite face of the hIL-9Rα binding site (**Fig. 2j**). The B-helix in the bound form is markedly more fixed and is moved away from the core by 4 Å, while maintaining its helicity and flexibility upon receptor engagement (**Fig. 2k**). A second event concerns the conformation of helix D, which in the unbound state sways away from the helical core of IL-9 by 5-10°. However, in the bound form the D-helix moves into a position even closer to the core of the cytokine than any state in the ensemble (**Fig. 2j**). Interestingly, γc-binding cytokines typically use the A- and D-helices to bind to γ_c_. Thus, binding of hIL-9Rα to the AC-face of the cytokine leads to conformational priming of IL-9 to enable recruitment of γ_c_ to the AD-face on the opposite face of the cytokine.

### The structure of mouse IL-9 reveals structural divergence in helix-B

Because the structure of hIL-9 revealed several distinct features, we deemed it opportune to compare it with mIL-9, an ortholog frequently used to study and model IL-9 function in physiology and disease. In order to achieve high-resolution structural information on mIL-9 we utilized the Fab fragment of the anti-mIL-9 antibody 35D8 to aid crystallization efforts of this cytokine. This anti-mIL-9 antibody was shown to bind mIL-9 with high affinity (**Fig. S5a**). The Fab:mIL-9 complex led to crystals that diffracted synchrotron X-rays to 1.8 Å resolution (**Table 2**). The Fab:mIL-9 complex reveals that Fab 35D8 binds mIL-9 with a polar interaction footprint over an interface area of 785 Å^2^ covering both the A- and C-helix of the cytokine (**Fig. S5b, Table S1**). The structure of mIL-9 confirms the conservation of the ten cysteines engaged in disulfide bonds (**Fig. S5b**). By superposing hIL-9 on mIL-9 we can detect some differences between the two orthologs next to their general similarity (r.m.s.d=0.7 Å for 547 of 797 atoms) (**Fig. S5c**). The most pronounced difference is the longer B-helix in mIL-9 compared to hIL-9 (**Fig. S5c**). This is paired with the loss of the helical turn in the BC-loop. This indicates that the B-helix is not a conserved structural feature among IL-9 orthologs, which suggest that it is not essential for the function of IL-9.

### The IL-9 system is distinct from receptor assemblies mediated by IL-2/IL-15 and IL-7

The fact that the B-helix is not conserved between hIL-9 and mIL-9 and therefore likely not essential in IL-9 signaling, indicates that there is no conserved binding site present for a possible third receptor, as has been suggested recently based on structure predictions of IL-9 (*27*). To situate IL-9 within the IL-2 cytokine family the trimmed helical cores of the cytokines, for which a structure was available, were aligned and subjected to a phylogenetic analysis (**Fig. S5d**). The resulting phylogenetic tree places IL-9 between the IL-7 and IL-4/IL-13 branches, away from the IL-2/IL-15 branches. This phylogenetic analysis together with the observation that the B-helix is not structurally conserved, indicates that the IL-9 signaling complex is most likely not like the IL-2/IL-15 system and that IL-9 is not missing a third receptor. Additionally, IL-9 has been linked to IL-7 by sequence similarity (26% sequence identity between mature hIL-7 and hIL-9) and similar gene structures, causing them to be classified as a subgroup within the family (*31*). In hindsight, a sequence identity of 26% and the striking presence of ten cysteines in the sequence of IL-9, while there are only six in IL-7, should have already alerted us to a substantially different structure of IL-9 compared to IL-7. However, with the structure of hIL-9 in unbound and bound states we can more fully interrogate this relationship (**Fig. S5e**). One of the more marked traits of IL-7 is the kink in its A-helix due to a π-helical turn, which is a feature it shares with TSLP (*32, 33*). From the determined structure of hIL-9 we can conclude that IL-9 does not share this feature as its A-helix is a standard α-helix (**Fig. S5e**). Additionally, the position of the AB-loop in respect to the D-helix has been shown to differ amongst cytokines, with the AB-loop of the long-chain cytokines running in front of the D-helix whereas for most of the short-chain cytokines the AB-loop runs atop of the D-helix (*34, 35*). IL-7 and TSLP are outliers of the IL-2 family with their AB-loop running in front of the D-helix, adding to their close relationship (*27*). Also, in this aspect IL-9 differs from IL-7 as the AB-loop in IL-9 runs atop of the D-helix (**Fig. S5e**). Additionally, the distinct short B-helix of IL-9 is also not represented in IL-7. The lengths of the four helices of IL-7 all fall within expectation, while the B-helix in IL-9 is evidently shorter. Together, these observations indicate that the structure of IL-9 does not agree with the classification of an IL-7-like fold. Therefore, our structural analysis indicates that IL-9 should be regarded as a separate branch of IL-7.

### hIL-9Rα is a structural outlier within the IL-2 cytokine receptor family

The structure of the binary IL-9:IL-9Rα complex also allows for a comparison between hIL-9Rα and the other hγ_c_-binding receptors, namely hIL-2Rβ, hIL-4Rα, hIL-7Rα and hIL-21Rα. When superposing these receptors onto hIL-2Rβ, two uniquely structural features of hIL-9Rα become apparent (**Fig. 3**). Firstly, the FG2-loop is extended in hIL-9Rα. A structure-based sequence alignment indicates that this extension is due to the insertion of a largely acidic sequence, which is absent in the other receptors (**Fig. 3b**). This extended loop is however clearly conserved in hIL-9Rα orthologs (**Fig. S6**). Part of this loop (Glu238-Val235) is buried upon interaction with the N-terminal part of hIL-9. This extended loop presents several potential hurdles for hγ_c_ to bind in a similar way to that described for the IL-2, IL-4, and IL-15 complexes. The part of the loop that is projected towards hγ_c_ is highly negatively charged (**Fig. 3c**), the position of the loop would sterically hinder the engagement of hγ_c_ as described for the other complexes and furthermore, if this partially ordered loop would become more ordered upon binding hγ_c_ this would increase the entropic penalty of this binding event. As a second specific feature of hIL-9Rα, we can distinguish a surface-exposed histidine following the β-strand D2 in the stem region (His200), whereas in the other receptors the β-strand continues to this position, which is generally a conserved hydrophobic residue (Ile/Val/Leu) (**Fig. 3d,e**). As this β-strand is part of the interaction site with hγ_c_ in the determined hγ_c_-complexes, a disruption with an outward-pointing histidine will result in a distinctly different binding interface and might have an impact on the binding of hγ_c_. It is clear from these features that hIL-9Rα will not be able to engage hγ_c_ in a similar way without making severe adjustments.

**Figure 3:**
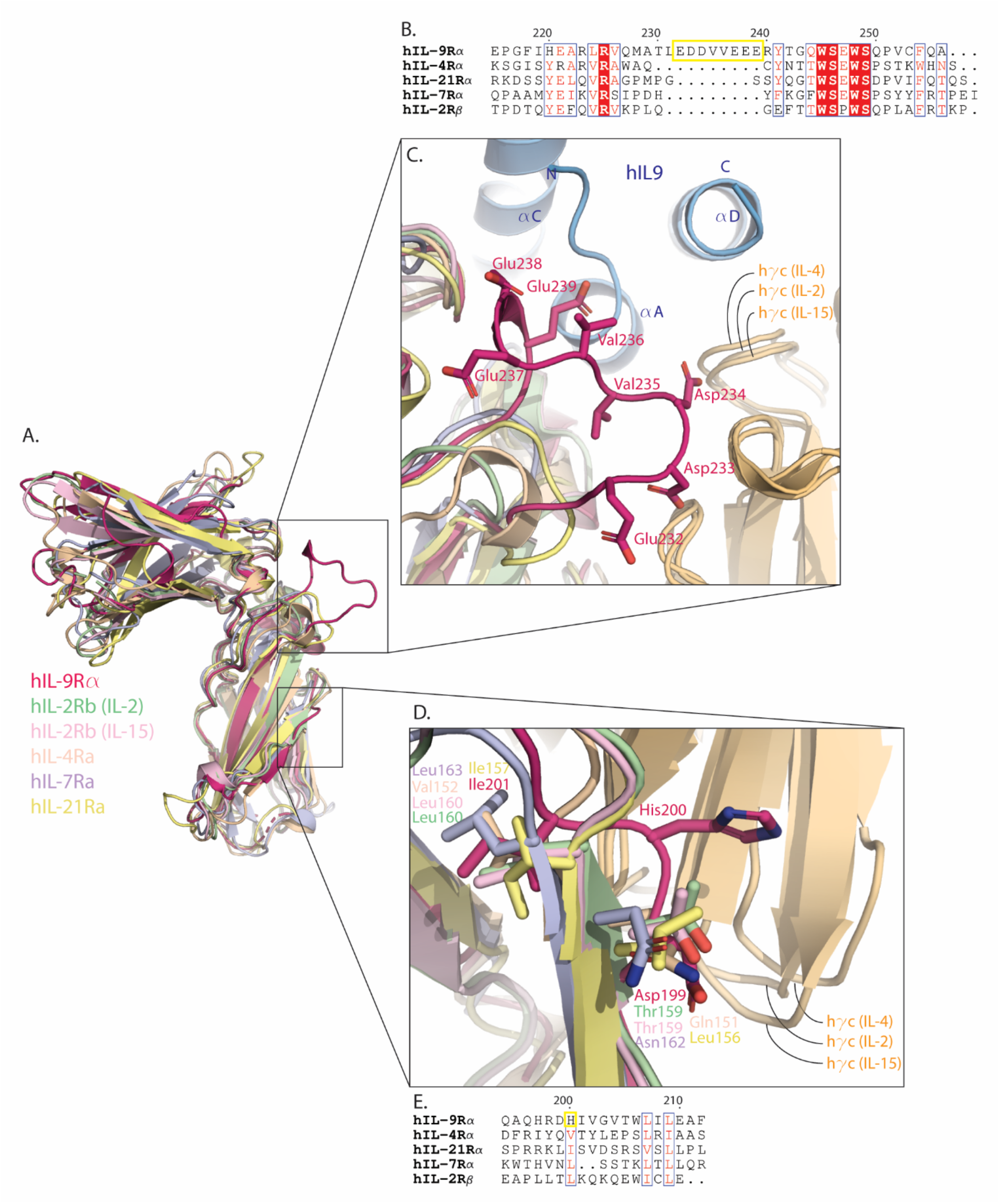
hIL-9Rα is a structural outlier among the receptors that pair with γc. (A) Structural superposition of all hγ_c_-interacting receptors, namely hIL-2Rβ from the hIL-2 complex, hIL-4Rα, hIL-7Rα, hIL-9Rα, hIL-2Rβ from the hIL-15 complex and hIL-21Rα. (B) Structure-based multiple sequence alignment of the γc-interacting receptors. Inserted acidic sequence in hIL-9Rα indicated in yellow. Conserved residues are shown on a red background. Semi-conserved residues are boxed and shown in red. (C) Close-up view of the FG2-loop of the γc-interacting receptors. Inserted sequence in hIL-9Rα is shown in sticks. hIL-9 is shown in blue, hγ_c_ from the hIL-2, hIL-4 and hIL-15 is shown in orange. (D) Close-up view of the region around the D2 β-strand of the γc-interacting receptors. Relevant residues are shown in sticks, hγc from the hIL-2, hIL-4 and hIL-15 is shown in orange. (E) Structure-based multiple sequence alignment of the γc-interacting receptors. His200 of hIL-9Rα is highlighted in yellow. Well conserved residues are boxed and shown in red.

### SNP rs2069885 in hIL-9 does not impact the kinetics or affinity of receptor binding

The single nucleotide polymorphism (SNP) rs2069885 resulting in a T117M point mutation in hIL-9 has been linked to asthma exacerbation, laryngeal squamous cell carcinoma (LSCC) and gender-specific effects on lung function, cystic fibrosis, and RSV infections in infants (*36*–*40*). This is the most prevalent missense SNP in hIL-9, with an allele frequency of 0.11 according to the genome aggregation database. Mapping of this point mutation to our structure of hIL-9, shows that this residue is situated distant from the hIL-9Rα interface, on the CD-loop as part of the anti-parallel β-strands (**Fig. S7**). To determine whether this mutation affects the kinetics of the signaling complex we performed additional BLI experiments with hIL-9^T117M^ (**Fig. S7**). The results show that there is no difference in the kinetics or affinity in the formation of the hIL-9^T117M^-mediated binary or ternary complex compared to those mediated by wild-type IL-9.

### Antagonism of hIL-9 bioactivity by monoclonal antibodies

With IL-9 involved in multiple inflammatory diseases, IL-9 has emerged as a potential drug target. In this context, monoclonal antibodies (mAbs) targeting hIL-9 were developed aimed at inhibition of the IL-9 signaling pathway. The three selected antibodies bind hIL-9 with high affinity with similar association and dissociation rates as determined by BLI (**Fig. 4a-c**). However, they show significantly different capacities in inhibiting the IL-9 signaling pathway in a cellular proliferation assay (**Fig. 4d**). The presence of antibody 6E2 resulted in the inhibition of IL-9-induced proliferation with an IC50 of 31 pM (**Fig. 4d**). Antibody 6D3 exerts its inhibitory function with an IC50 of 58 pM and antibody 7D6 performs poorly in comparison, with an IC50 of 2.1 nM.

**Figure 4:**
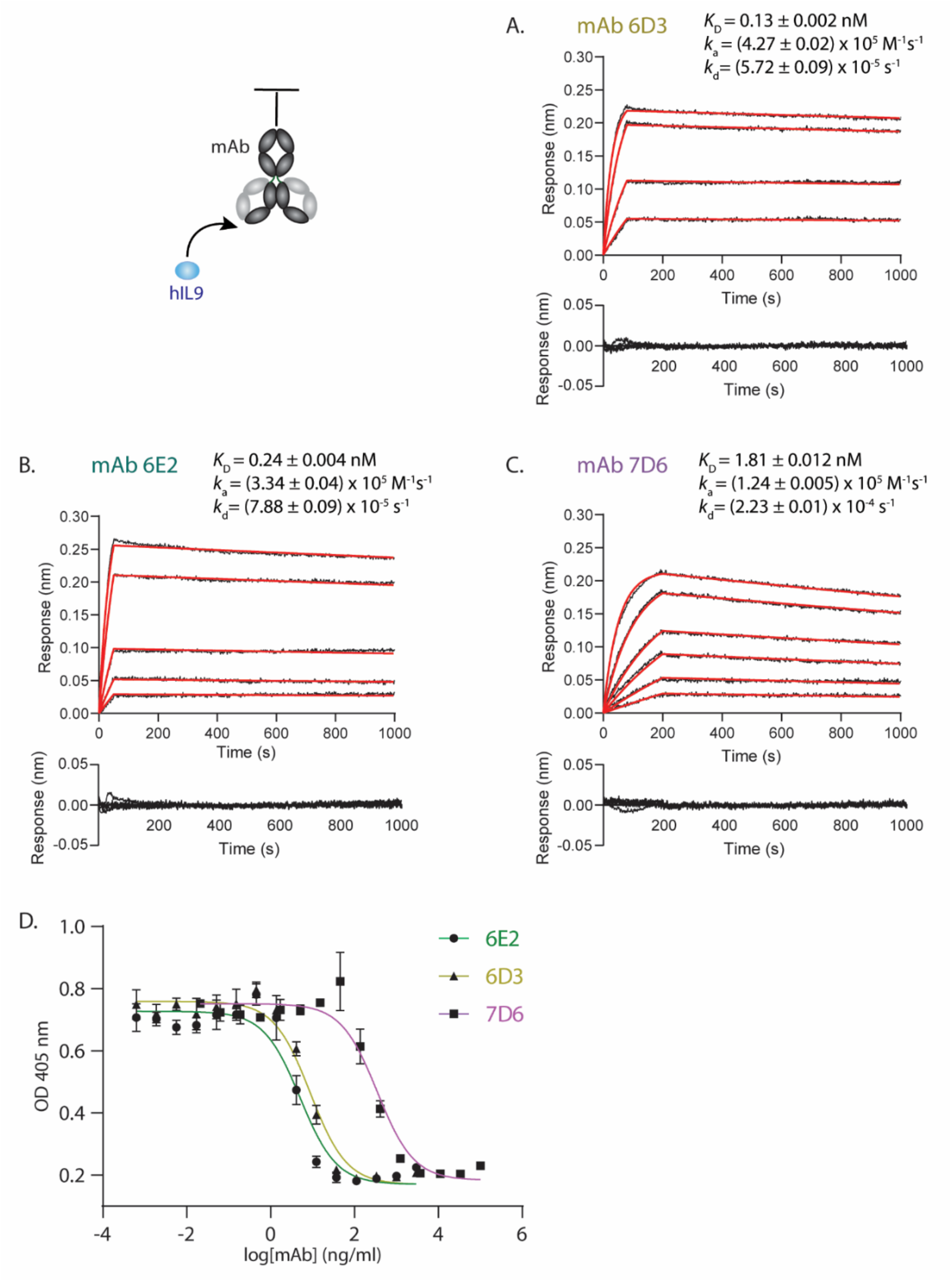
Antagonistic antibodies against human IL-9. (A,B,C) Representative BLI data traces of the binding of hIL-9 (two-fold dilution series, lowest concentration of 5 nM) to mAb coupled to AHC sensors. Data traces were fitted with a 1:1 model with residuals shown below. (A) For mAb 6D3. (B) For mAb 6E2. (C) For mAb 7D6. (D) Proliferation assay using Ba/F3-h9R cells stimulated with hIL-9 and increasing concentrations of mAb (6D3, 6E2 or 7D6). IC50 values for 6E2: 31 pM, for 6D3: 58 pM, for 7D6: 2.1 nM.

### The mAbs binding epitopes overlap with the hIL-9Rα binding site

To gain structural insights and to elucidate the antagonistic properties of the anti-hIL-9 antibodies, we determined the corresponding Fab:hIL-9 complexes by X-ray crystallography. The Fab:hIL-9 complexes were biochemically reconstituted (**Fig. S1**) and set up for crystallization. Hits were optimized and data was collected for well-diffracting crystals. The first Fab:hIL-9 complex (Fab 6D3:hIL-9) crystal resulted in a dataset to 1.7 Å resolution (**Table 2**). The electron density map after crystallographic phasing by molecular replacement using a model for the Fab, allowed us to build hIL-9 de novo preventing model biases. In terms of the Fab:hIL-9 interface, Fab 6D3 targets hIL-9 by mainly binding the C-helix and the first half of the A-helix with a polar footprint covering an interface area of 750 Å^2^ (**Fig. 5a, Table S1**). Two Arg-Asp interactions govern the specificity of the interaction (**Fig. 5a**). This includes Arg91 in hIL-9, which is also involved in the interaction with hIL-9Ra. Next, two additional Fab:hIL-9 complexes were structurally determined (**Fig. 5b,c and Table 2**). Fab 6E2 binds hIL-9 with an average interface area of 987 Å^2^. The cytokine is placed with its A-helix in-between the light and heavy chain of the Fab (**Fig. 5b**) (**Table S1**). Hereby, the light chain interacts specifically with the C-helix by coordinating Arg91 of the cytokine through Asp31 and Asp49 (**Fig. 5b**). Another Asp on the light chain, Asp95, interacts with the main chain of Leu24 placed at the tip of helix A. The heavy chain of Fab 6E2 adds to this interaction site by engaging with the A-helix and part of the D-helix. Next, Fab 7D6 mainly binds the A-helix, with the heavy chain covering the first half of the helix and the light chain the second half creating an interface area of 862.8 Å^2^ (**Fig. 5c and Table S1**). Even though the interaction mainly covers the A-helix, the light chain still engages Arg91 through an interaction with the main chain of Lys94 on the light chain (Fig 5c). Thus, in all Fab:hIL-9 complexes Arg91 of the cytokine is engaged in a specific interaction. Next, to understand how the antibodies inhibit the binding of hIL-9 to its receptors and how we can explain the different efficacies of the three antibodies in inhibiting the IL-9 signaling pathway, we superposed the structure of the binary hIL-9:hIL-9Ra complex with the structure of the Fab:hIL-9 complexes (**Fig. 5d**). Our analysis reveals that the binding epitopes of all these Fabs, and therefore the corresponding mAbs, (partially) overlap with the binding site of hIL-9Rα on IL-9. Thus the developed Fabs are competitive receptor antagonists.

**Figure 5:**
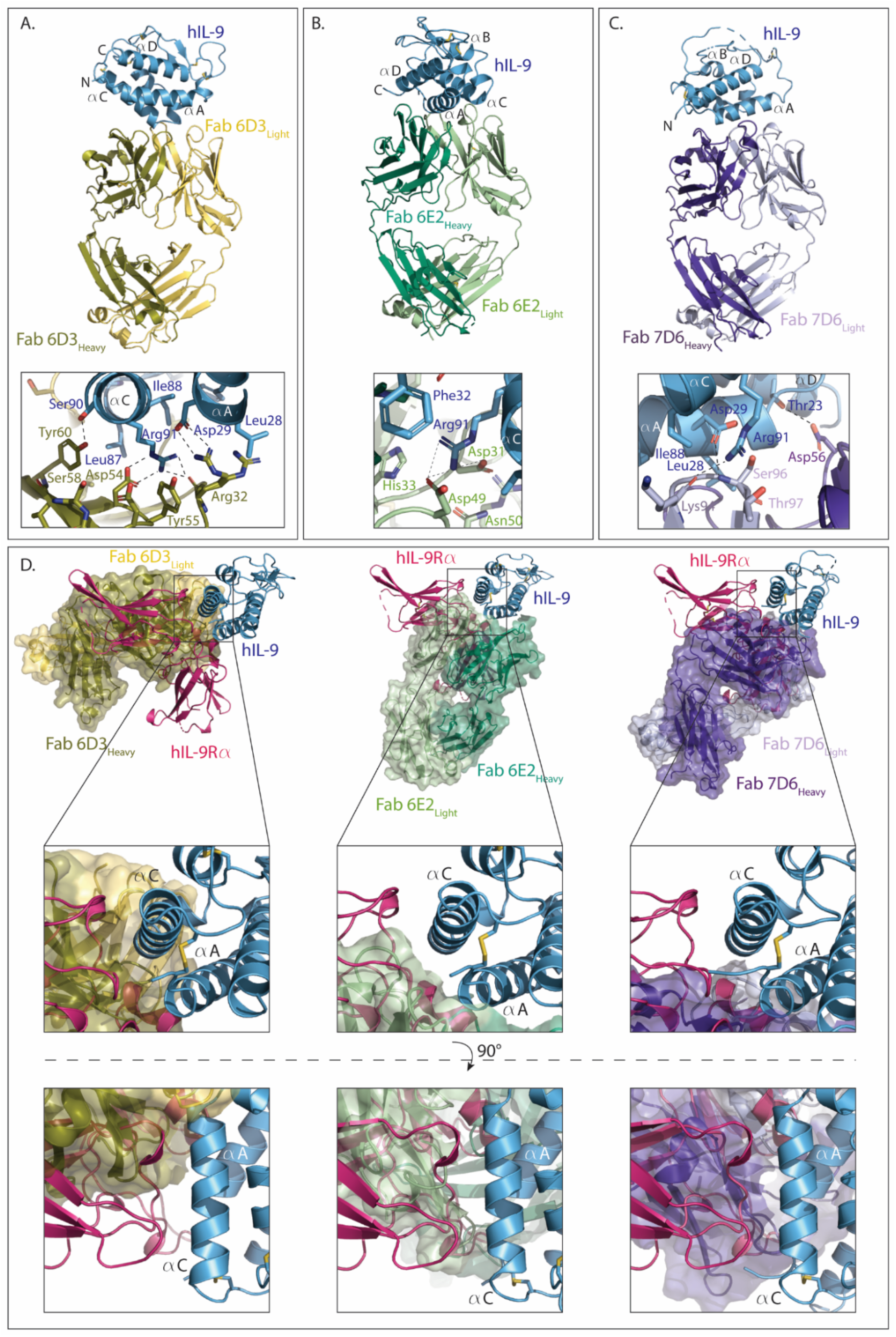
Structural basis for antagonism of hIL-9 by anti-IL-9 antibodies. (A,B,C) Cartoon representation of Fab:hIL-9 complexes with a close-up view of the interaction in the vicinity of Arg91 of the cytokine below (A) for the Fab 6D3:hIL-9 complex, (B) for the Fab 6E2:hIL-9 complex and (C) the Fab 7D6:hIL-9 complex. (D) Overlay of the Fab:hIL-9 complexes with the hIL-9:hIL-9Rα complex based on the structural superposition of hIL-9 with a close-up front view and top view of the binding of the C-helix by the Fabs.

### Helix C in hIL-9 guides binding to hIL-9Rα

Antibody 7D6 performs poorly in inhibiting hIL-9 in the cellular proliferation assay whereas 6E2 performs very well (IC50 = 2.1 nM versus 31 pM) (**Fig. 4d**). Since the large difference in potency cannot be simply explained by the small difference in affinity (*K*_D_ = 1.8 nM versus 0.24 nM) (**Fig. 4a-c**) we took a closer look at the epitopes of these Fabs. Both Fab fragments cover the A-helix extensively. However, their orientations on hIL-9 are perpendicular to each other. This results in Fab 6E2 holding hIL-9 in a crevice between its light and heavy chains, covering also the C-helix to a large extent (**Fig. 5b,d**). Fab 7D6, on the other hand, hardly covers the C-helix (**Fig. 5c,d**). This availability of the C-helix might create an entry point for hIL-9Rα to engage hIL-9 through the C-helix and force the dissociation of the Fab from hIL-9. Next, we bring Fab 6D3 into this comparison. This antibody performs only slightly worse than antibody 6E2 in inhibiting the hIL-9 signal in the cellular reporter assay (IC50 = 58 pM versus 31 pM) (**Fig. 4d**). The structure of Fab 6D3:hIL-9 shows us that it binds mainly through the C-helix and leaves the A-helix almost completely available (**Fig. 5a,d**). This indicates that availability of the A-helix is not enough to allow hIL-9Rα to efficiently bind and dislodge the Fab. When examining the epitope on the C-helix, we see that Fab 6D3 does not completely cover the hIL-9Rα binding site, which could explain the difference in efficacy between mAb 6E2 and mAb 6D3 (**Fig. 5d**). This indicates that the efficacy of the antibodies in inhibiting the hIL-9:hIL-9Rα interaction and the hIL-9 signaling pathway, is correlated with the ability of the antibodies to cover the hIL-9Rα binding site on the C-helix of hIL-9. This shows that the C-helix is key mediator of the interaction between hIL-9 and hIL-9Rα.

## Discussion

Interleukin-9, important in Th9 and ILC2 immunity, has been linked to a multitude of inflammatory diseases, autoimmune diseases, and cancer types. It is the last member of the IL-2 cytokine family for which no structural and biophysical data describing its association with receptors are available. The findings presented here resolve this critical structure-function bottleneck.

We show that hIL-9 binds hIL-9Rα with a low nanomolar affinity in an entropy-driven binding mode, which is reflected in the largely hydrophobic cytokine-receptor interface. By dissecting the structures of hIL-9 in complex with anti-hIL-9 Fabs and the corresponding efficacies of the antibodies in inhibiting hIL-9 signaling, we show that the C-helix is the main entry point for hIL-9Rα on hIL-9. We also show that the mode of action of these antibodies was by sterically hindering the binding of hIL-9Rα to hIL-9. In addition, we show that the hIL-9:hIL-9Rα interaction leads to a restructuring of the B- and D-helix in hIL-9, where the D-helix is fixed to a position closer to the helical core. As the other γ_c_-binding cytokines use the AD-face to engage with γc, this might be a necessity for the binding of hγ_c_ by hIL-9 as there is no measurable affinity between hIL-9 and hγ_c_ without hIL-9Rα present. Even with these structural rearrangements, the resulting affinity of hγc for the binary complex is weak, with a micromolar affinity partially caused by a drastically fast dissociation rate. This low affinity of the ternary complex, measured by ITC and BLI, is in accordance with an early report where the binding of mIL-9 on MC/9 cells (mouse mast cell line) was unchanged when mγ_c_ was blocked by an antibody, indicating that the binding of mIL-9 to the cells is solely dependent on mIL-9Rα (*41*).

A possible explanation for the low affinity of the ternary complex is provided by the comparison of hIL-9Rα with the other hγ_c_-engaging receptors where two features of hIL-9Rα stand out. Firstly, hIL-9Rα contains an inserted acidic sequence in the FG2-loop, which might several possible difficulties for hγc to bind. As this inserted sequence is negatively charged, hγ_c_ will need to present complemetnary charges on its side. In addition, comparison with other receptors pairing with hγ_c_ shows that this loop in IL-9Rα is substantially longer, indicating that it might sterically impact the approach of hγ_c_. Lastly, if this loop were to become ordered in the ternary complex, it would present a large entropic penalty since this loop still retains some flexibility in the binary hIL-9:hIL-9Rα complex. Secondly, hIL-9Rα contains a histidine where other receptors place a hydrophobic residue as part of the D2 β-strand. As the residues of this β-strand are involved in the interaction with hγc in the hIL-2, hIL-4 and hIL-15 complexes, this outward-pointing histidine will have a direct effect on this region of the binding interface. In conclusion, hIL-9Rα is a structural outlier among the hγc-binding receptors, potentially complicating the engagement with hγ_c_ with similarly high affinities to related cytokine-receptor assemblies.

This raises the question of why the IL-9-mediated extracellular complex is so unstable. A possible explanation might be to keep γ_c_ available as a shared receptor for the other members of the IL-2 cytokine family that signal through γ_c_. It was shown that the availability of γ_c_ represents a level of regulation within this family (*42, 43*). In that regard, the high affinity of the binary complex might function as a way to keep IL-9 bound on the cell surface, ready to engage γc when the availability is high enough.

We have shown that hIL-9 is structurally different from hIL-7, despite their similar sequences and gene structures, as it does not resemble the key features of hIL-7, being the kinked A-helix and the AB-loop running in front of the D-helix. This is also valid vice-versa, where the features of hIL-9, including the shortened B-helix, are not represented in hIL-7. It has recently been proposed that the IL-9 signaling complex may be more similar to IL-2 and IL-15 and that it might be missing a third receptor (*27*). We successfully utilized the Fab fragment of an anti-mIL-9 antibody to aid in crystallization of mIL-9, which allowed us to compare the two orthologs. This reveals that the B-helix is not structurally conserved, despite a large overall similarity in sequence. This allows us to speculate that there is no conserved binding site in this region and that consequently there would not be a third receptor for IL-9. This also becomes apparent in the phylogenetic analysis where the IL-9 branch is placed away from the IL-2/IL-15 branch.

We also investigated a prevalent SNP (rs2069885) that results in a point mutation T117M in hIL-9, which is associated with gender-specific outcomes in lung diseases. We produced hIL-9^T117M^ recombinantly in mammalian cells and subjected it to additional biophysical studies. We could not determine a difference in kinetics or affinity in the formation of the binary or ternary complex. Additionally, we mapped the mutation on the structure, indicating that the affected residue is far from the binding site of hIL-9Rα and, based on homologous structures, predicted to be away from the binding site for hγ_c_. How this point mutation would affect disease outcomes, will require additional investigation.

## Methods

### Production of recombinant hIL-9, mIL-9 and hIL-9Rα in *E. coli*

Codon-optimized cDNA fragments (Ginkgo Bioworks) encoding for mature hIL-9 (NP_000581.1, residues 19-144), mature mIL-9 (NP_032399.1, residues 19-144) and the ectodomain of hIL-9Rα (NP_002177.2, residues 40-261) were cloned in the pET15b vector in frame with a N-terminal hexahistidine tag followed by a thrombin cleavage site. hIL-9, mIL-9 and hIL-9Rα were expressed by IPTG induction in *E. coli* BL21(DE3) strain at 310 K as inclusion bodies and refolded in vitro based on an in-house protocol (*29*). Following refolding, the protein is captured via immobilized metal affinity chromatography (IMAC) and purified by size exclusion chromatography (SEC). The N-terminal tag was removed by an overnight incubation with thrombin (Merck). Undigested protein was removed by IMAC. The flowthrough containing the protein of interest was polished via SEC with HBS pH 7.4 as running buffer.

### Production of ^13^C,^15^N isotopically labelled hIL-9 in *E. coli*

Isotopically labelled hIL-9 was produced in *E. coli* BL21(DE3) cells transformed with the pET15b-hIL-9 expression construct as described above. Cells were grown in minimal medium at 310 K supplemented with ^15^N-ammonium chloride and ^13^C-D-glucose. Purification was performed as described above with the final SEC using 20mM NaH_2_PO_4_, pH 6.5, 100mM NaCl as running buffer.

### Production of hIL-9, hIL-9^T117M^ and hγc^23-262^ in mammalian cells and hγc in insect cells

The codon-optimized cDNA fragment for mature hIL-9 (NP_000581.1, residues 19-144) and the codon-optimized cDNA fragment for the complete extracellular domain of hγc^23-262^ (NP_000197.1, residues 23-262) were cloned with a chicken RTPμ-like signal peptide and a C-terminal caspase-3 site followed by an AviHis-tag. The T117M point mutation was introduced in hIL-9 (hIL-9^T117M^) by overlap extension mutagenesis and cloned in the same vector. Transient expression was performed in adherent HEK293T cells using polyethylenimine as transfection agent (*44*). Cells were maintained in DMEM supplemented with 10% fetal calf serum (FCS). Prior to transfection, the medium was exchanged for DMEM supplemented with 3.6 mM valproic acid. Conditioned medium was harvested by centrifugation after 4 days and filtered through a 0.22 μm filter before proceeding to the chromatographic steps.

The cDNA fragment for the trimmed extracellular domain of hγc (NP_000197.1, residues 56-254) was cloned with a gp64 signal peptide and a C-terminal caspase-3 site followed by an AviHis-tag. Expression was achieved using the baculovirus system. *Spodoptera frugiperda* (Sf9) cells were grown at 301K in Insect-XPRESS medium (Lonza Bioscience). First, high titers of virus were generated. This virus was then added to the Sf9 cells in log phase to produce recombinant protein. The conditioned medium was harvested after 5 days by centrifugation and filtered through a 0.22 μm filter before proceding to the chromatographic steps.

The protein of interest was captured from the clarified conditioned medium by IMAC using a cOmplete His-Tag purification column (Roche) and further purified by SEC using preparative grade HiLoad 16/600 Superdex 75 columns (Cytiva) with HBS pH 7.4 as running buffer. To remove the C-terminal tags, the protein was incubated overnight with caspase-3. Undigested protein was removed by IMAC. The flowthrough containing the protein of interest was polished via SEC with HBS pH 7.4 as running buffer.

### Llama immunization and library construction

Llamas were immunized intramuscularly once per week over six weeks with carrier free recombinant human IL-9 or mouse IL-9 (R&D Systems). Each llama (4 in total) received 40 μg of IL-9, buffered in phosphate-buffered saline (PBS) and mixed with Incomplete Freund’s Adjuvant (Sigma-Aldrich), the first two weeks, and 20 μg the remaining four weeks. Generation of Ab libraries was performed using the SIMPLE antibody™ platform at argenx as previously described (*45*). Briefly, five days after the last immunization, peripheral blood lymphocytes were purified and used for extraction of total RNA. Total RNA was then converted into random primed cDNA using reverse transcriptase, and gene sequences encoding for VH-CH1 regions of llama IgG and VL-CL domains (kappa and lambda) were isolated by PCR and subcloned into a phagemid vector pCB3.

### Phage selection for Fab generation

The *E. coli* strain TG1 (Netherlands Culture Collection of Bacteria) was transformed using recombinant phagemids to generate Fab-expressing phage libraries (one lambda and one kappa library per immunized llama). The phages were adsorbed on immobilized recombinant biotinylated IL-9, and eluted using trypsin as previously described (*46*). Three rounds of phage display selections were performed to enrich for phages expressing IL-9-specific Fabs. TG1 *E. coli* was finally infected with selected phages, and individual colonies were isolated. Secretion of Fabs was induced using isopropyl β-D-1-thiogalactopyranoside (Sigma-Aldrich), and the Fab-containing periplasmic fractions of bacteria were collected and screened by SPR using a Biacore 3000 apparatus (GE Healthcare).

### Monospecific Ab production, purification and characterization

The cDNAs encoding the VH and VL (lambda or kappa) domains of the neutralizing Fabs fragments displaying the lowest off-rate were cloned into two separate mammalian expression vectors (U-Protein Express BV) which comprise the cDNAs encoding the CH1, hinge, CH2 and CH3 domains of a human IgG1 Ab (or mouse IgG2a), containing a mutation that abrogates Ab effector functions mediated by the Fc receptor, or the CL (lambda or kappa), respectively. Production by transient transfection of HEK293 cells and endotoxin-free purification by protein A affinity chromatography was then performed to generate human IgG1 (or mouse IgG2a) monoclonal Abs. Their purity and homogeneity were verified by SDS-PAGE (2 μg of each sample), and their high affinity were determined by SPR. The neutralizing activity of anti-IL-9 monospecific Abs was finally confirmed in in vitro cellular assays, in which human IL-9 induced the proliferation of Ba/F3-h9R cells. For crystallographic studies, the cDNAs encoding the VHs were recloned into a vector containing only the CH1. After transfection of HEK293 cells of this plasmid with the light chain plasmid (same for the full monoclonal production), the Fabs were purified from the supernatant using Capture Select IgG-CH1 Affinity beads (Life Technologies) according to the manufacturer indications.

### NMR structure determination of hIL-9

All NMR spectra were recorded at CEITEC Josef Dadok National NMR Centre on 850 MHz Bruker Avance III spectrometer equipped with ^1^H/^13^C/^15^N TCI cryogenic probe head with z-axis gradients. Three sparsely sampled 4D NMR experiments were acquired: 4D HC(CC-TOCSY(CO))NH, 4D ^13^C,^15^N edited HMQC-NOESY-HSQC (HCNH), and 4D ^13^C,^13^C edited HMQC-NOESY-HSQC (HCCH). Sequential and aliphatic side chain assignments were obtained automatically using the 4D-CHAINS algorithm (*30*). Analysis of the chemical shifts of all ten cysteines in hIL9 reflected an oxidized state of the attached sulfur atoms. The cysteine Cα were upfield (range 52.5-56.5 p.p.m.) and Cβ downfield (range 39.0-42.5 p.p.m.) shifted consistent with the presence of five intramolecular disulfide bridges. A first calculation was exclusively based on NOE-derived distance constraints from the two 4D NOESY spectra (HCNH and HCCH). Inspection of the assigned NOE crosspeaks revealed H^β^-H^β^ NOE correlations between cysteines that allowed to identify unambiguously three cysteine bridges. In a second calculation the three cysteine pairs were linked and the remaining pairs were deduced based on proximity in the resulting structures. In a last calculation all cysteine pairs were linked and an automated NOE cross-peak assignment was performed using the software CYANA 3.0 (*47*). The distance restraints from the CYANA calculation, TALOS-derived dihedral angle restraints (*48*), and disulfide restraints were used in a water refinement calculation (*49*) applying the RECOORD protocol (*50*). The quality of the NMR-derived structure ensembles was validated using PSVS (*51*).

### Crystal structure of hIL-9:hIL-9Rα complex

The binary complex was formed by adding a molar excess of refolded hIL-9 to refolded hIL-9Rα. The binary complex was isolated and separated from the molar excess hIL-9 by SEC using a Superdex 75 column with HBS pH 7.4 as running buffer. Fractions containing the binary complex were pooled and concentrated to 8.5 mg/ml. Commercial sparse matrix screens were set up using a Mosquito liquid handling robot (TTP Labtech) in a vapor-diffusion sitting drop setup with 100nl of mother liquid mixed with 100nl protein solution in SwissSci 96-well triple drop plates. Plates were incubated at 293K. A hit was observed in the ProPlex screen (Molecular Dimensions) condition A7 (0.1M ammonium sulfate, 0.1 M TRIS, pH 7.5, 20% PEG1500). The condition was optimized with the best-diffracting crystals grown in 0.1 M ammonium sulfate, 0.1 M TRIS, pH 5.6, 18% PEG 1500. Crystals were cryoprotected with mother liquor supplemented with 20% ethylene glycol prior to being cryo-cooled in liquid nitrogen. Diffraction data was collected at 100 K at the PROXIMA 2A beamline at SOLEIL, Paris.

Diffraction data was integrated using the XDS suite (*52*). Initial phases were determined by maximum-likelihood molecular replacement (MR) as implemented in the program suite PHASER (*53*) using hIL-9 from the crystal structure of the Fab 6D3:hIL-9 complex and IL-2Rβ (PDB: 2B5I) as a model for hIL-9Rα. A pair of hIL-9 models in a relative orientation, a solution from a previous molecular replacement run, needed to be placed first in order to successfully place the IL2Rβ model. There are 8 copies in the asymmetric unit. Model building and refinement were iteratively performed in COOT (*54*), PHENIX (*55*) and AutoBuster (*56*). NCS restraints, reference model restraints (using hIL-9 from the crystal structure Fab 6D3:hIL-9 complex) and riding hydrogen model were applied during refinement. Refinement was performed for individual coordinates and atomic displacement parameters (ADP). Model and map validation tools in COOT, PHENIX and PDB_REDO server (*57*) were used throughout the work flow to guide the process.

### Crystal structures of Fab:h/mIL-9 complexes

Fab:h/mIL-9 complexes were formed by adding a molar excess of refolded h/mIL-9 to the Fab fragment. The Fab:h/mIL-9 complex was separated from the molar excess h/mIL-9 by SEC with HBS pH 7.4 as running buffer. Fractions containing the Fab:h/mIL-9 complex were pooled and concentrated. Commercial sparse matrix screens were set up using a Mosquito liquid handling robot (TTP Labtech) in a vapor-diffusion sitting drop setup with 100nl of mother liquid mixed with 100nl protein solution in SwissSci 96-well triple drop plates. Plates were incubated at 293K.

The Fab 6D3:hIL-9 complex was concentrated to 14 mg/ml. A hit was observed in the ProPlex screen condition A7. Final optimized condition with the best-diffracting crystals was 0.1 M ammonium sulfate, 0.1 M Tris pH 5.8, 16% PEG 1500. Crystals were protected with mother liquor supplemented with PEG 1500 increased to 25% prior to being cryo-cooled in liquid nitrogen. Diffraction data was collected at 100 K at the P14 beamline at PETRA 3, Hamburg. Diffraction data was integrated using the XDS suite (*52*). Initial phases were determined by MR as implemented in the program suite PHASER (*53*). Trimmed models for the variable domain and constant domain of the Fab were made using PDB 5NIV chain B as a model for the heavy chain and PDB 6CT7 chain B as a model for the light chain. The structure of hIL-9 was built de novo in the electron density. There is 1 copy in the asymmetric unit. Model building and refinement were iteratively performed in COOT (*54*) and PHENIX (*55*). Refinement was performed for individual coordinates and ADP combined with a translation-libration-screw rotation (TLS) model. Model and map validation tools in COOT, PHENIX and PDB_REDO server (*57*) were used throughout the work flow to guide the process.

The Fab 6E2:hIL-9 complex was concentrated to 13.5 mg/ml. A hit was observed in the Wizard screen (Molecular Dimensions) condition D3. The final optimized condition with the best-diffracting crystals was 0.2 M lithium sulfate, 0.1 M phosphate/citric acid pH 3.8, 21% PEG 1000. Crystals were protected with mother liquor supplemented with 20% glycerol prior to being cryo-cooled in liquid nitrogen. Diffraction data was collected at 100 K at the ID23-1 beamline at ESRF, Grenoble. Diffraction data was integrated using the XDS suite (*52*). Initial phases were determined by MR using the program suite PHASER (*53*). Trimmed models for the variable domain and constant domain of the Fab were made using PDB 6GHG chain A as model for the heavy chain and PDB 5Y9J chain L as model for the light chain. As model for hIL-9 a poly-ALA structure of hIL-9 as determined in the crystal structure for the Fab 6D3:hIL-9 complex was used. There are 4 copies in the asymmetric unit. Model building and refinement were iteratively performed in COOT (*54*), PHENIX (*55*) and AutoBuster (*56*). Refinement was performed for individual coordinates and ADP. NCS restraints, reference model restraints (using hIL-9 from the crystal structure Fab 6D3:hIL-9 complex) and riding hydrogen model were applied during refinement. Model and map validation tools in COOT, PHENIX and PDB_REDO server (*57*) were used throughout the work flow to guide the process.

The Fab 7D6:hIL-9 complex was concentrated to 15 mg/ml. A hit was observed in the PEG/Ion screen (Hampton Research) condition B7 and the BCS screen (Molecular Dimensions) condition D3. The final optimized condition with the best-diffracting crystals was 0.25 M ammonium nitrate, 18% PEG 3350, 5% ethylene glycol with a 2:1 ratio (protein:mother liquor). Crystals were protected with 0.25 M ammonium nitrate, 22% PEG 3350, 20% ethylene glycol prior to being cryo-cooled in liquid nitrogen. Diffraction data was collected at 100 K at the MASSIF-3 beamline at ESRF, Grenoble. Diffraction data was integrated using the XDS suite (*52*). Initial phases were determined by MR using the program suite PHASER(*53*). Trimmed models for the Fab were made using PDB 4HC1 chain H as model for the heavy chain and PDB 4R90 chain L as model for the light chain. The structure of hIL-9 was built de novo in the electron density. There are 4 copies in the asymmetric unit. Model building and refinement were iteratively performed in COOT (*54*), PHENIX (*55*) and AutoBuster (*56*). Refinement was performed for individual coordinates and ADP. NCS restraints and riding hydrogen model were applied during refinement. Model and map validation tools in COOT, PHENIX and PDB_REDO server(*57*) were used throughout the work flow to guide the process.

The Fab 35D8:mIL-9 complex was concentrated to 13.5 mg/ml. A hit was observed in the PEG/Ion screen (Hampton Research) condition C2. The final optimized condition with the best-diffracting crystals was 0.25 M zinc acetate, 23% PEG 3350 with a ratio 1:1.6 ratio (protein:mother liquor). Crystals were protected with mother liquor supplemented with 20% glycerol/DMSO/ethylene glycol (in 1:2:2 ratio) prior to being cryo-cooled in liquid nitrogen. Diffraction data was collected at 100 K at the P13 beamline at PETRA 3, Hamburg. Diffraction data was integrated using the XDS suite (*52*). Initial phases were determined by MR as implemented in the program suite PHASER (*53*). Trimmed models for the variable and constant domain of the Fab were made using PDB 4ZS7 (chain H) as model for the heavy chain and PDB 5BV7 chain L as model for the light chain. The structure of mIL-9 was built de novo in the electron density. There is 1 copy in the asymmetric unit. Model building and refinement were iteratively performed in COOT (*54*) and PHENIX (*55*). Refinement was performed for individual coordinates and ADP combined with TLS parameterization. The riding hydrogen model was applied during refinement. Model and map validation tools in COOT, PHENIX and PDB_REDO server (*57*) were used throughout the work flow to guide the process.

### In vitro biotinylation of recombinant hIL-9 and hγc

In vitro biotinylation of glycosylated hIL-9, hIL-9^T117M^, hγc and hyc^23-262^ was achieved by incubating the protein carrying the Avi-tag with a 1/100 molar ratio of BirA biotin ligase (*58*), 3mM biotin and 5mM ATP overnight at 293K. The excess biotin is removed by a desalting step using the HiPrep 26/10 Desalting column (Cytiva). The biotinylated protein was aliquoted and flashfrozen in liquid nitrogen.

### Biolayer interferometry

BLI experiments on the IL-9 signaling complex were performed using an Octet Red 96 machine (Sartorius) in kinetics buffer (20 mM HEPES pH 7.4, 300 mM NaCl, 0.05% (w/v) BSA, 0.05% (v/v) Tween 20) at 298 K. Streptavidin (SA) sensors (Sartorius) were functionalized with in vitro biotinylated ligands and quenched with a biotin solution. The functionalized tips were then dipped in different analyte concentrations. Non-functionalized tips were used as negative controls in a double referenced setup. After subtracting the control sensorgrams, the resulting data was fitted with a 1:1 binding model. Data analysis was performed using the Data Analysis software 9.0.0.14 (Sartorius).

BLI experiments on the antagonistic antibodies were performed using an Octet Red 96 machine (Sartorius) in kinetics buffer (PBS, 0.1% (w/v) BSA, 0.02% (v/v) Tween20) at 298 K. Anti-hIgG Fc capture (AHC) or anti-mIgG Fc capture sensors (Sartorius) were functionalized with monoclonal antibodies. The functionalized tips were then dipped in different hIL-9 or mIL-9 (both from R&D systems) concentrations. Non-functionalized tips were used as negative controls in a double referenced setup. After subtracting the control sensorgrams, the resulting data was fitted with a 1:1 binding model. Data analysis was performed using the Data Analysis software 9.0.0.14 (Sartorius).

### Isothermal Titration Calorimetry

Prior to all measurements all protein pairs were buffer matched on SEC in standard SEC running buffer (20 mM HEPES pH 7.4, 150 mM NaCl) unless mentioned otherwise. Experiments were carried out using a MicroCal PEAQ-ITC instrument. The experiments were conducted at 310 K or 298 K. Peak analysis was performed using NITPIC and data analysis was performed using SEDPHAT with an AB hetero-association model while allowing the syringe concentration to be corrected for concentration errors and inactive fractions.

### Structure-based multiple sequence alignment of hγc-binding receptors

Multiple sequence alignment was performed using PROMALS3D in standard setting for hIL-9Rα, hIL-2Rβ (PDB: 2B5I), hIL-4Rα (PDB: 3BPL), hIL-7Rα (PDB: 3DI2) and hIL-21Rα (PDB: 3TGX) and in a separate analysis for all structurally determined mouse and human cytokines of the IL-2 cytokine family, using only the 4 helices of the four-helical bundle without loops and termini. This includes hIL-2 (PDB: 2B5I), mIL-2 (PDB: 4YUE), hIL-4 (PDB: 3BPL), hIL-7 (PDB: 3DI2), hIL-13 (PDB: 3BPO), hIL-15 (PDB: 4GS7), mIL-15 (PDB: 2PSM), hIL-21 (PDB: 3TGX), hTSLP (PDB: 5J11), mTSLP (PDB: 4NN6) together with hIL-9 and mIL-9.

### Phylogenetic analysis of cytokines

To analyze the relation between the cytokines, we generated a phylogenetic tree upon the multiple sequence alignment using MrBayes with the following parameters. We set the priors to a fixed “wag” amino acid rate matrix model (prset aamodelpr=fixed(wag)), the likelihood model to a gamma distribution with invariable sites and 4 categories (lset rates=invgamma Ngammacat=4), and carried Markov chain Monte Carlo (MCMC) simulations over five runs of 500,000 generations, with sampling frequency of 10 (mcmc nchains=4 ngen=500000 nruns=5 printfreq=1000 samplefreq=10 savebrlens=yes starttree=random). Branch support was evaluated by posterior probabilities after a burn-in of 25%. After the simulations the topological convergence diagnostic was lower than 0.005. The phylogenetic tree was generated using the iTOL server.

### Cell culture

MO7e cells were obtained from Leibnitz Institute, German Collection of Microorganisms and cell cultures GmbH (DSMZ) (code: ACC 104) and cultured in RPMI (Gibco) medium supplemented with 10 ng/ml human GM-CSF and 20% FCS (Gibco). For the purposes of proliferation assays, cytokine starvation was done for 6h in RPMI and 5% FCS.

### Bioactivity assay hIL-9

MO7e cells were washed two times with RPMI, cytokine starved for 6h and divided into 96-well plate (20 000 cells/well). Stimulation was done using increasing concentrations of hIL-9 and the cells were cultivated for 90 h at 310 K and 5% CO_2_. The total volume in each well was 100 μl, and all the stimulations were done in triplicates. Cells treated with GM-CSF served as the positive control. To measure the cytokine-dependent proliferation cells were incubated for 15 min with 50 μl of Promega CellTiter-Glo Viability assay 2.0 solution. The signal was measured in the GloMax Luminometer using 1s integration time, values were normalized to untreated cells and relative proliferation calculated. The data were plotted and fitted to a log agonist versus response curve in GraphPad Prism® software and the EC50 was determined.

### Competition binding assay

Inhibitory activity of hIL-9 antibodies was measured on Ba/F3-h9R cells that respond to hIL-9. The cloning of hIL-9Rα and stable transfection of Ba/F3 cells was previously described (*5*). Ba/F3-h9R cells were maintained in Dulbecco’s modified Eagle medium (DMEM) containing 10% fetal calf serum and IL-3 (250 U/ml, produced in CHO transfected cells). 40U/ml of hIL-9 was pre-incubated with indicated concentration of different anti-hIL-9 antibodies for 30 minutes. Ba/F3-h9R cells were washed three times with DMEM and 3000 Ba/F3-h9R cells were added per well. Cells were incubated at 310 K, 8% CO2 for 3 days, and proliferation was measured by hexosaminidase activity determination. Substrate of hexosaminidase was added 2h30 before the measurement.

## Supplementary materials

**Fig. S1:**
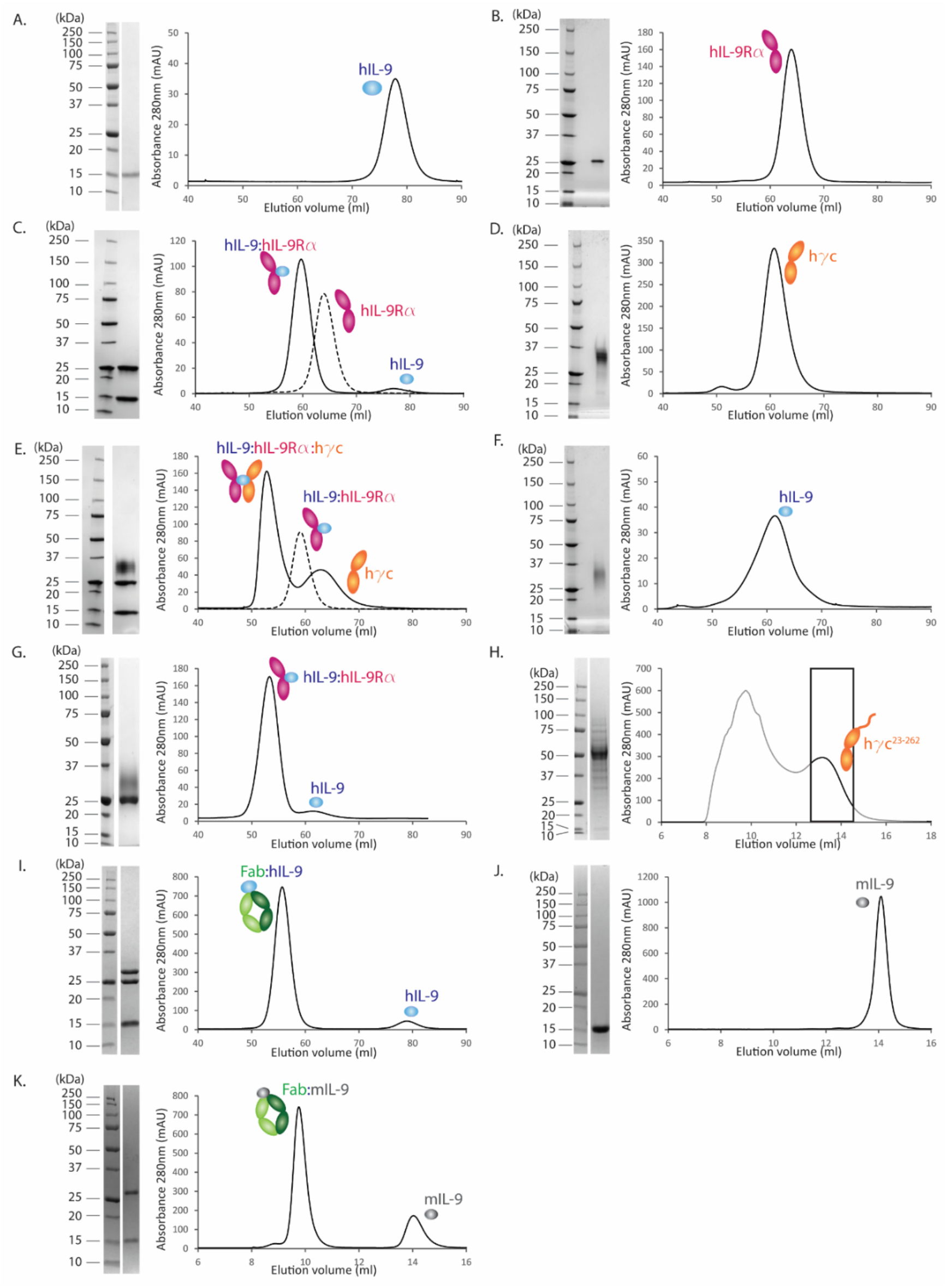
Purification and biochemical reconstitution of IL-9-mediated complexes. SDS-PAGE analysis and SEC elution profiles on a HiLoad 16/600 Superdex 75 pg column for (A) hIL-9 produced in *E. coli*, (B) hIL-9Rα produced in *E. coli*, (C) binary hIL-9:hIL-9Rα complex with excess hIL-9 (solid line) and hIL-9Rα (dashed line), (D) hγc produced in Sf9 cells, (E) ternary hIL-9:hIL-9Rα:hγc complex with excess hγc and binary hIL-9:hIL-9Rα complex (dashed line), (F) glycosylated hIL-9 produced in HEK293T cells, (G) binary hIL-9:hIL-9Rα complex with excess glycosylated hIL-9. (H) SDS-PAGE analysis and SEC elution profile on a Superdex 200 Increase 10/300 GL column for hyc^23-262^ produced in HEK293T cells. (I) Representative SDS-PAGE analysis and SEC elution profile on a HiLoad 16/600 Superdex 75 pg column for a Fab:hIL-9 complex with excess non-glycosylated hIL-9. (J) SDS-PAGE analysis and SEC elution profiles on a Superdex 75 Increase 10/300 GL column for mIL-9 produced in *E. coli* and (K) for Fab 35D8:mIL-9 complex with excess of mIL-9.

**Fig. S2:**
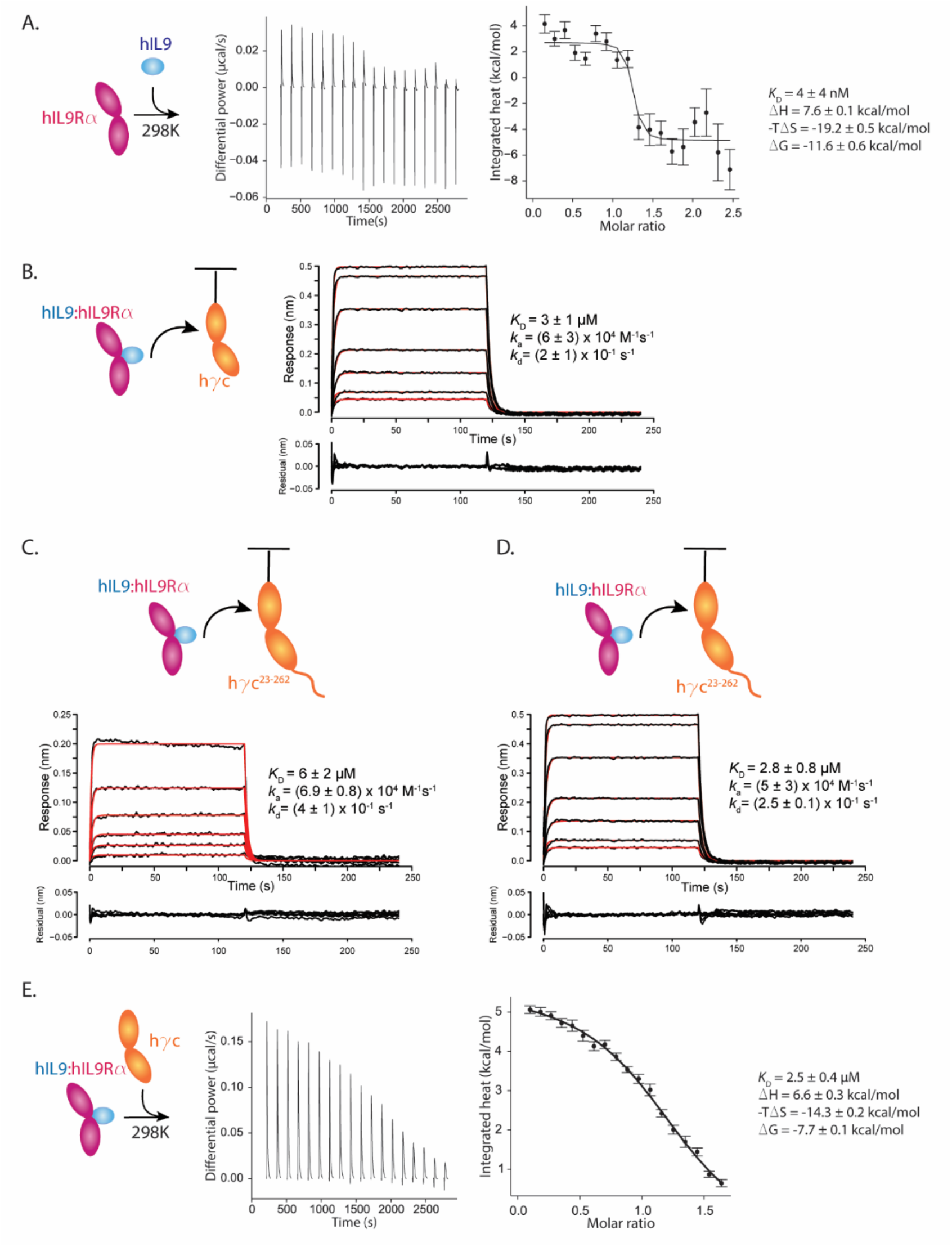
Additional characterization of the binary and ternary complex by BLI and ITC. (A,B,C) Representative BLI experiments were performed using the Octet RED96 at 298 K. Data traces were fitted with a 1:1 model with residuals shown below. (A) Binding of non-glycoysylated binary hIL-9:hIL-9Rα complex (highest concentration of 10 μM, two-fold dilution series) to coupled hγc. The represented values are averaged values and their s.d. from six replicate experiments. (B) Binding of binary hIL-9:hIL-9Rα complex containing glycosylated hIL-9 (highest concentration of 7.5 μM, two-fold dilution series) to coupled hγc^23-262^. The represented values are averaged values and their s.d. from three replicate experiments. (C) Binding of non-glycosylated binary hIL-9:hIL-9Rα complex (highest concentration of 10 μM, two-fold dilution series) to coupled hγc^23-262^. The represented values are averaged values and their s.d. from three replicate experiments. (D,E) Representative ITC experiments performed using a PEAQ-ITC instrument at 298 K. (D) Titration of 20.6 μM hIL-9 in 1.6 μM hIL-9Rα at 298 K with a local correction factor of 0.84. The represented values are averaged values and their s.d. from two replicate experiments. (E) Titration of 266 μM hγc in 31 μM hIL-9:hIL-9Rα complex at 298 K with a local correction factor of 0.76. The represented values are averaged values and their s.d. from three replicate experiments.

**Fig. S3:**
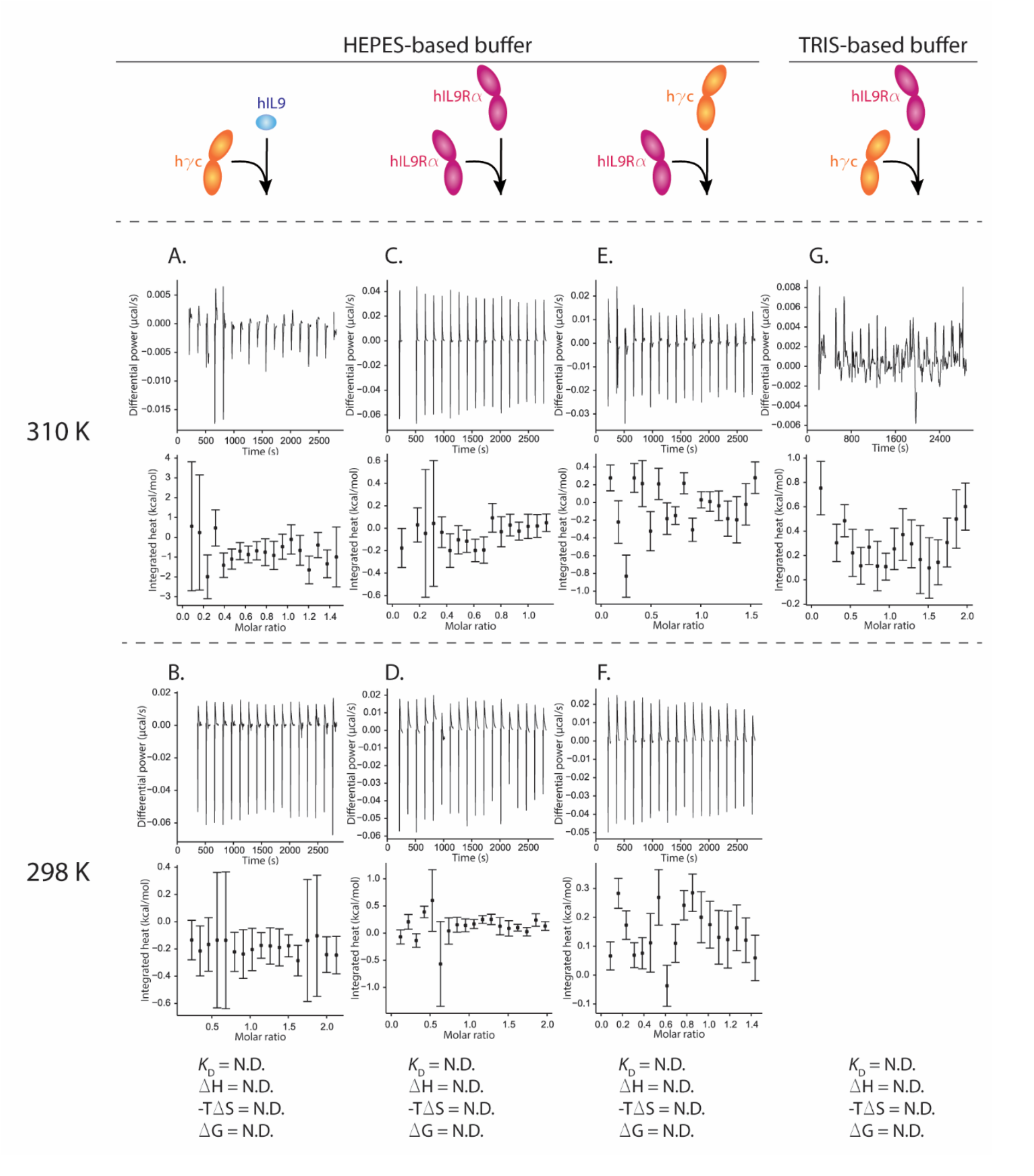
Thermodynamic characterization of interactions between various components of the hIL-9 signaling complex at 298 K and 310 K. Representative ITC experiments performed using a PEAQ-ITC instrument at 310 K (upper half) or at 298 K (lower half) in buffers as indicated on top. HEPES-based buffer contains 20mM HEPES, 150 mM NaCl, pH 7.4. TRIS-based buffer contains 20mM TRIS, 150mM NaCl, pH 7.4. (A) Titration of 151 μM hIL-9 in 19.7 μM hγc at 310 K. (B) Titration of 247 μM hIL-9 in 22.2 μM hγc at 298K. (C) Titration of 168 μM hIL-9Rα in 28.3 μM hIL-9Rα at 310 K. (D) Titration of 252 μM hIL-9Rα in 24.4 μM hIL-9Rα at 298 K. (E) Titration of 181 μM hγc in 22.4 μM hIL-9Rα at 310 K. (F) Titration of 100 μM hγc in 9.6 μM hIL-9Rα. (G) Titration of 175 μM hIL-9Rα in 16.9 μM hγc at 310 K.

**Fig. S4:**
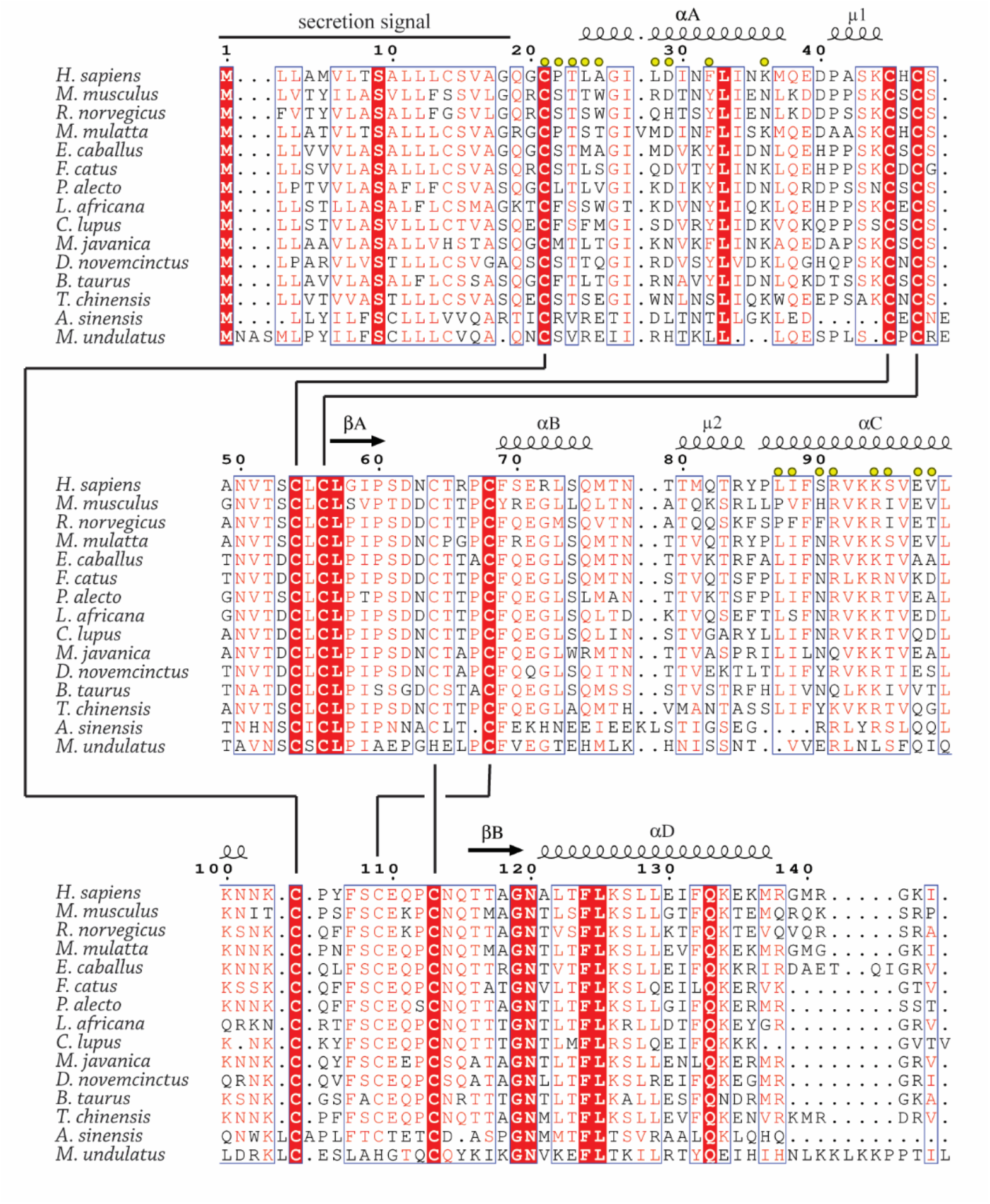
Sequence alignment of IL-9 orthologs. Sequence alignment of selected IL-9 orthologs. Residue numbering and structural annotations correspond to human IL-9. Conserved residues are shown on a red background. Disulphide pairs are connected by black lines. Semi-conserved residues are boxed and shown in red. Yellow spheres indicate residues interacting with hIL-9Rα. Multiple sequence alignment was performed using the MAFFT server (https://www.ebi.ac.uk/Tools/msa/mafft/) and annotated using the ESPripT server (http://espript.ibcp.fr/ESPript/ESPript/index.php).

**Fig. S5:**
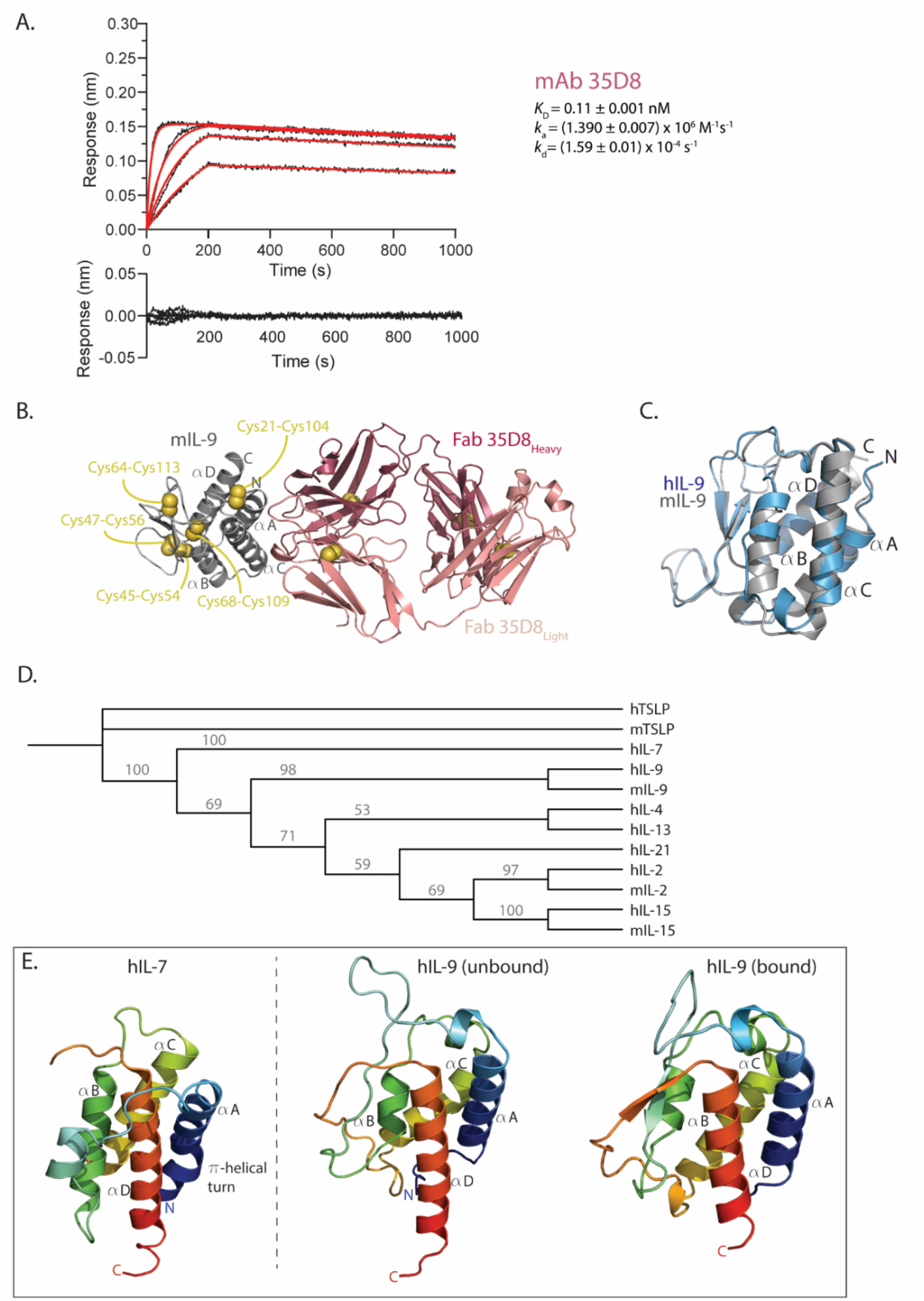
IL-9 in the IL-2 cytokine family. (A) Representative BLI experiment and residuals of the binding of mIL-9 (highest concentration of 12.5 nM, two-fold dilution series) to coupled mAb 35D8 on AMC sensors. Data traces were fitted with a 1:1 model with residuals shown below. (B) Cartoon representation of the Fab 35D8:mIL-9 complex. (C) Structural superposition of hIL-9 (blue) and mIL-9 (grey). (D) Phylogenetic analysis of the trimmed helical bundles of the IL-2 family cytokines for which a structure is available. Confidence level are indicated on the branches in grey. (E) Side-by-side comparison of hIL-7 (PDB: 3DI2 chain A) and hIL-9 (unbound state and bound state) in cartoon representation. Blue-red color gradient from N- to C-terminus.

**Fig. S6:**
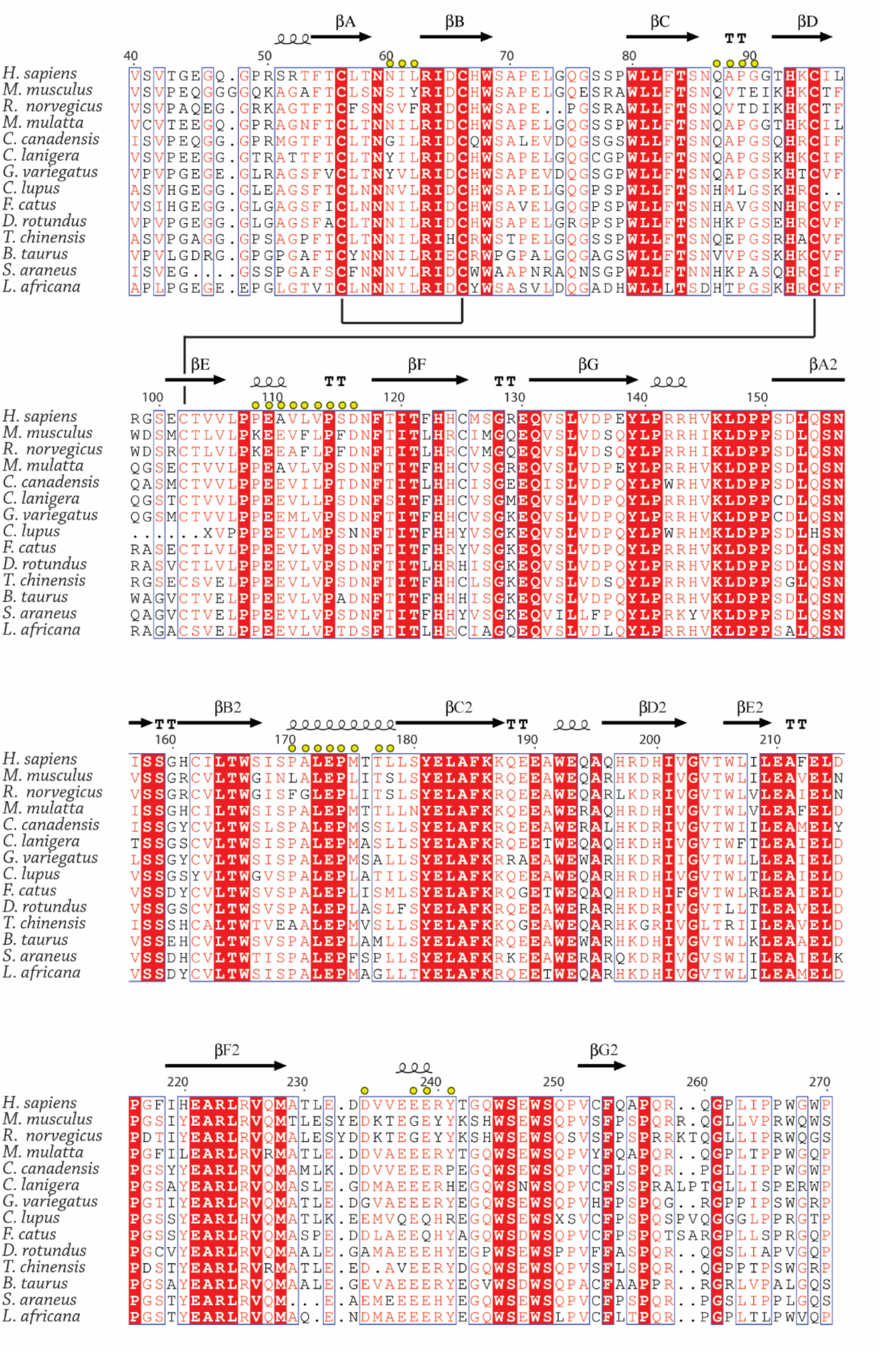
Sequence alignment of IL-9Rα orthologs. (A) Sequence alignment of selected IL-9Rα orthologs. Residue numbering and structural annotations correspond to human IL-9Rα. Conserved residues are shown on a red background. Disulphide pairs are connected by black lines. Semi-conserved residues are boxed and shown in red. Yellow spheres indicate residues interacting with hIL-9. The multiple sequence alignment was performed using the MAFFT server (https://www.ebi.ac.uk/Tools/msa/mafft/) and annotated using the ESPripT server (http://espript.ibcp.fr/ESPript/ESPript/index.php)(*59*).

**Fig. S7:**
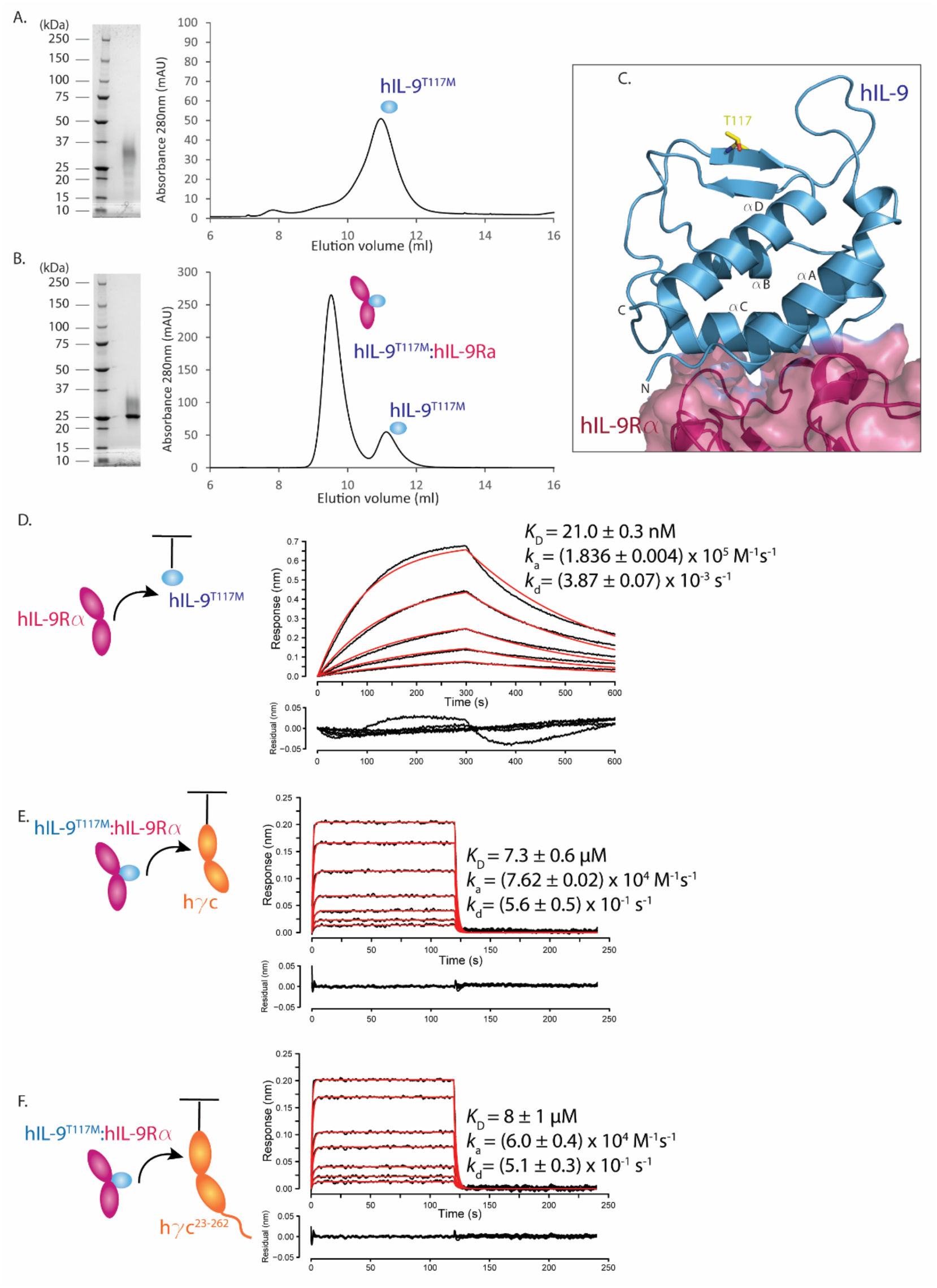
The point mutation T117M in hIL-9 has no apparent effect on the assembly and kinetics of receptor binding. (A) SDS-PAGE analysis and SEC elution profile on a Superdex 75 Increase 10/300 GL column for hIL-9^T117M^ produced in HEK293T cells. (B) SDS-PAGE analysis and SEC elution profile on a Superdex 75 Increase 10/300 GL column for hIL-9^T117M^:hIL-9Rα with an excess of hIL-9^T117M^. (C) Cartoon representation of hIL-9 with part of hIL-9Rα (in surface representation) to indicate the hIL-9:hIL-9Rα interface. Thr117 is indicated in yellow and represented as sticks. (D) Representative data traces and fit with corresponding residuals for the binding of hIL-9Rα (highest concentration of 140 nM, two-fold dilution series) to coupled hIL-9^T117M^. The represented values are averaged values and their s.d. from two replicate experiments. (E) Representative data traces and fit with corresponding residuals for the binding of hIL-9^T117M^:hIL-9Rα (highest concentration of 10.8 μM, two-fold dilution series) to coupled hγc. The represented values are averaged values and their s.d. from two replicate experiments. (F) Representative data traces and fit with corresponding residuals for the binding of hIL-9^T117M^:hIL-9Rα (highest concentration of 10.8 μM, two-fold dilution series) to coupled hγc^23-262^. The represented values are averaged values and their s.d. from two replicate experiments.

**Table S1:**
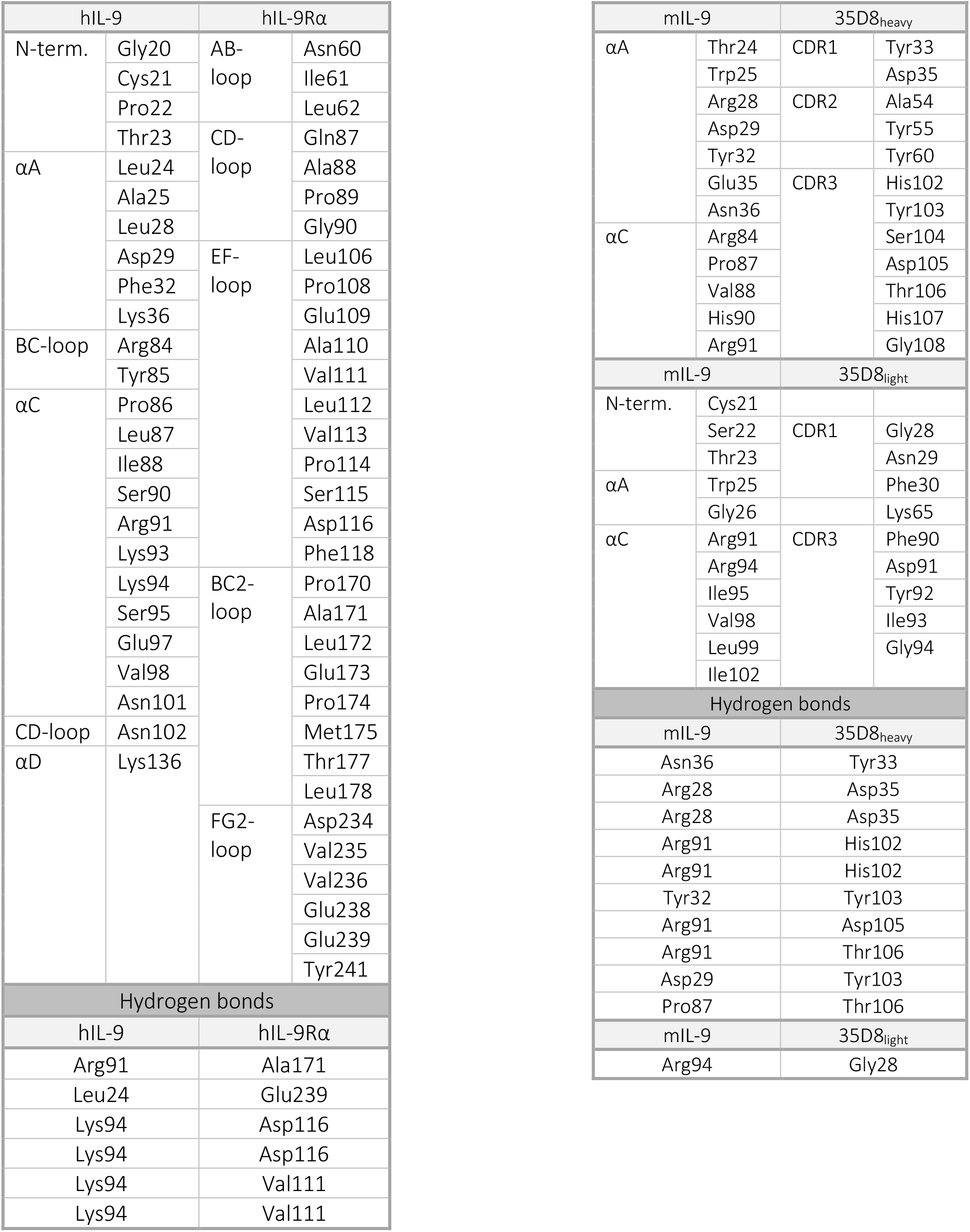
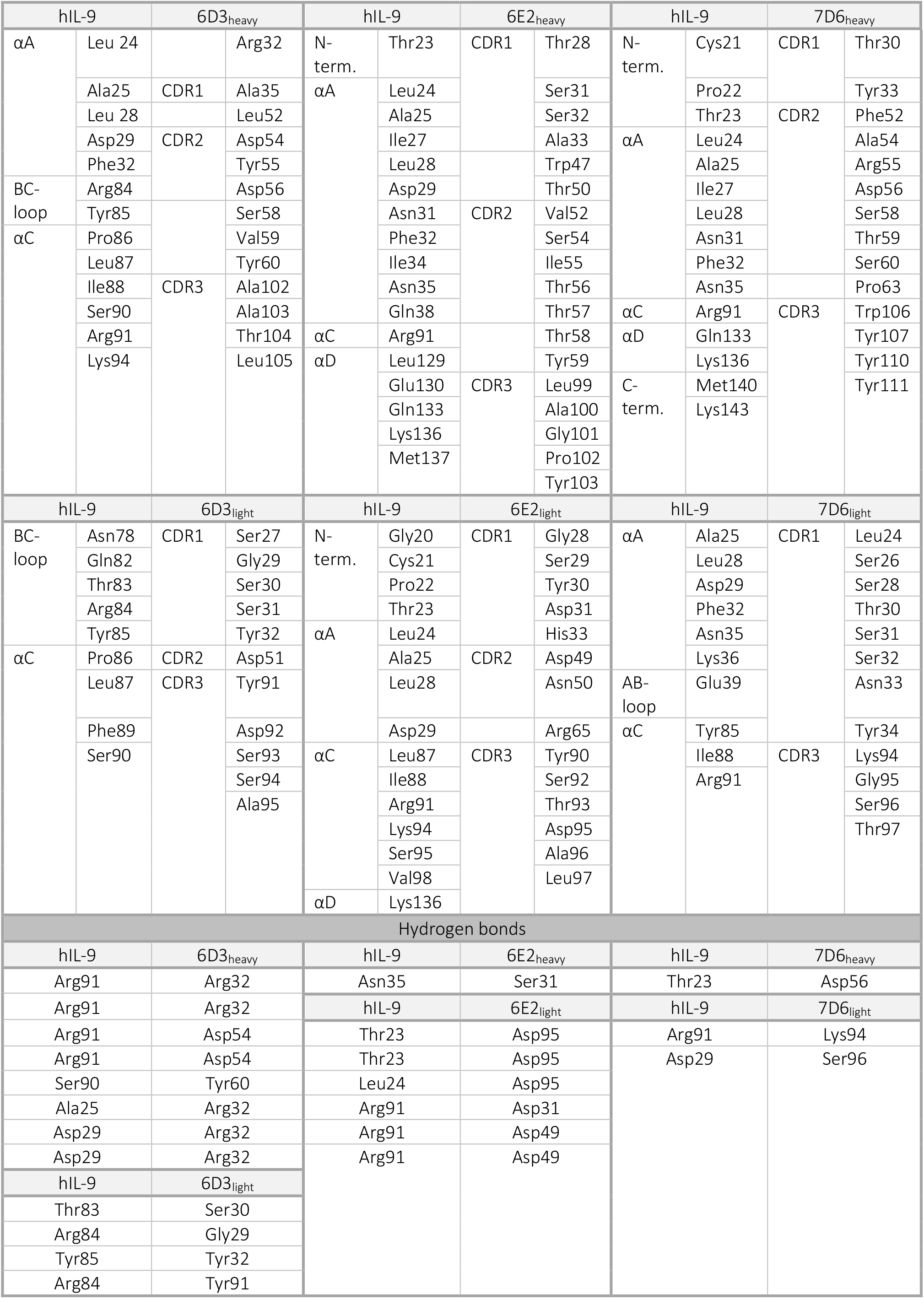
Overview of buried residues and hydrogen bonds at interaction interfaces. Buried residues and hydrogen bonds of interaction interfaces as analyzed by PISA. For the Fab:h/mIL-9 complexes, only the interfaces between Fab and h/mIL-9 were analyzed.

## Author contributions

T.D.V. carried out recombinant protein production of all IL-9 complex components, binding studies, structural studies by X-ray crystallography, and structural analyses with contributions from S.N.S. M.G. and F.B generated, characterized, and produced anti-IL-9 antibodies and Fab fragments thereof under supervision of C.B. I.M. performed the IL-9 proliferation assays. A.C.P. recorded and analyzed NMR spectra. T.E. assisted with NMR structure calculations. K.T. supervised the structural studies by NMR. E.M. performed the data processing and initiated refinement on the Fab6E2:hIL-9 crystal structure. L.D. performed cellular inhibition assays with the antibodies. T.D.V. and S.N.S wrote the manuscript with input from all authors. S.N.S. conceived and supervised the project.

## Data and materials availability

All structural coordinates reported in this work have been deposited in the Protein Data Bank (www.rcsb.org).

## References

1. C. Uyttenhove, R. J. Simpson, S. J. Van, Functional and structural characterization of P40, a mouse glycoprotein with T-cell growth factor activity. Proc.Natl.Acad.Sci.U.S.A. 85, 6934–6938 (1988).

2. L. Hültner, J. Moeller, E. Schmitt, G. Jäger, G. Reisbach, J. Ring, P. Dörmer, Thiol-sensitive mast cell lines derived from mouse bone marrow respond to a mast cell growth-enhancing activity different from both IL-3 and IL-4. J. Immunol. 142, 3440–6 (1989).

3. J. Moeller, L. Hültner, E. Schmitt, P. Dörmer, Partial purification of a mast cell growth-enhancing activity and its separation from IL-3 and IL-4. J. Immunol. 142, 3447–51 (1989).

4. L. Hültner, C. Druez, C. Uyttenhove, E. Schmitt, E. Rüde, P. Dörmer, J. Van Snick, Mast cell growth-enhancing activity (MEA) is structurally related and functionally identical to the novel mouse T cell growth factor P40/TCGFIII (interleukin 9). Eur. J. Biochem., 1413–1416 (1990).

5. J. B. Demoulin, C. Uyttenhove, E. Van Roost, B. DeLestré, D. Donckers, J. Van Snick, J. C. Renauld, A single tyrosine of the interleukin-9 (IL-9) receptor is required for STAT activation, antiapoptotic activity, and growth regulation by IL-9. Mol. Cell. Biol. 16, 4710–6 (1996).

6. J. B. Demoulin, J. Louahed, L. Dumoutier, M. Stevens, J. C. Renauld, MAP kinase activation by interleukin-9 in lymphoid and mast cell lines. Oncogene. 22, 1763–1770 (2003).

7. J. B. Demoulin, L. Grasso, J. M. Atkins, M. Stevens, J. Louahed, R. C. Levitt, N. C. Nicolaides, J. C. Renauld, Role of insulin receptor substrate-2 in interleukin-9-dependent proliferation. FEBS Lett. 482, 200–204 (2000).

8. V. Dardalhon, A. Awasthi, H. Kwon, G. Galileos, W. Gao, R. a Sobel, M. Mitsdoerffer, T. B. Strom, W. Elyaman, I.-C. Ho, S. Khoury, M. Oukka, V. K. Kuchroo, IL-4 inhibits TGF-beta-induced Foxp3+ T cells and, together with TGF-beta, generates IL-9+ IL-10+ Foxp3(-) effector T cells. Nat. Immunol. 9, 1347–1355 (2008).

9. M. Veldhoen, C. Uyttenhove, J. van Snick, H. Helmby, A. Westendorf, J. Buer, B. Martin, C. Wilhelm, B. Stockinger, Transforming growth factor-beta “reprograms” the differentiation of T helper 2 cells and promotes an interleukin 9-producing subset. Nat. Immunol. 9, 1341–1346 (2008).

10. C. Wilhelm, K. Hirota, B. Stieglitz, J. Van Snick, M. Tolaini, K. Lahl, T. Sparwasser, H. Helmby, B. Stockinger, An IL-9 fate reporter demonstrates the induction of an innate IL-9 response in lung inflammation. Nat. Immunol. 12, 1071–1077 (2011).

11. E. C. Nowak, C. T. Weaver, H. Turner, S. Begum-Haque, B. Becher, B. Schreiner, A. J. Coyle, L. H. Kasper, R. J. Noelle, IL-9 as a mediator of Th17-driven inflammatory disease. J. Exp. Med. 206, 1653–60 (2009).

12. Z. Wiener, A. Falus, S. Toth, IL-9 increases the expression of several cytokines in activated mast cells, while the IL-9-induced IL-9 production is inhibited in mast cells of histamine-free transgenic mice. Cytokine. 26, 122–130 (2004).

13. M. Stassen, M. Arnold, L. Hültner, C. Müller, C. Neudörfl, T. Reineke, E. Schmitt, Murine Bone Marrow-Derived Mast Cells as Potent Producers of IL-9: Costimulatory Function of IL-10 and kit Ligand in the Presence of IL-1. J. Immunol. 164, 5549–5555 (2000).

14. M. Xiao, Y. Wang, C. Tao, Z. Wang, J. Yang, Z. Chen, Z. Zou, M. Li, A. Liu, C. Jia, B. Huang, B. Yan, P. Lai, C. Ding, D. Cai, G. Xiao, Y. Jiang, X. Bai, Osteoblasts support megakaryopoiesis through production of interleukin-9. Blood. 129, 3196–3209 (2017).

15. B. R. Lauwerys, N. Garot, J.-C. Renauld, F. A. Houssiau, Cytokine Production and Killer Activity of NK/T-NK Cells Derived with IL-2, IL-15, or the Combination of IL-12 and IL-18. J. Immunol. 165, 1847–1853 (2000).

16. S. Takatsuka, H. Yamada, K. Haniuda, H. Saruwatari, M. Ichihashi, J. C. Renauld, D. Kitamura, IL-9 receptor signaling in memory B cells regulates humoral recall responses. Nat. Immunol. 19, 1025–1034 (2018).

17. D. S. Postma, E. R. Bleecker, P. J. Amelung, K. J. Holroyd, J. Xu, C. I. M. Panhuysen, D. A. Meyers, R. C. Levitt, Genetic Susceptibility to Asthma — Bronchial Hyperresponsiveness Coinherited with a Major Gene for Atopy. N. Engl. J. Med. 333, 894–900 (1995).

18. N. C. Nicolaides, K. J. Holroyd, S. L. Ewart, S. M. Eleff, M. B. Kiser, C. R. Dragwa, C. D. Sullivan, L. Grasso, L. Y. Zhang, C. J. Messler, T. Zhou, S. R. Kleeberger, K. H. Buetow, R. C. Levitt, Interleukin 9: a candidate gene for asthma. Proc. Natl. Acad. Sci. U. S. A. 94, 13175–80 (1997).

19. U. A. Temann, G. P. Geba, J. A. Rankin, R. A. Flavell, Expression of interleukin 9 in the lungs of transgenic mice causes airway inflammation, mast cell hyperplasia, and bronchial hyperresponsiveness. J. Exp. Med. 188, 1307–1320 (1998).

20. J. Kearley, J. S. Erjefalt, C. Andersson, E. Benjamin, C. P. Jones, A. Robichaud, S. Pegorier, Y. Brewah, T. J. Burwell, L. Bjermer, P. A. Kiener, R. Kolbeck, C. M. Lloyd, A. J. Coyle, A. A. Humbles, IL-9 governs allergen-induced mast cell numbers in the lung and chronic remodeling of the airways. Am. J. Respir. Crit. Care Med. 183, 865–75 (2011).

21. G. Seumois, C. Ramírez-Suástegui, B. J. Schmiedel, S. Liang, B. Peters, A. Sette, P. Vijayanand, Single-cell transcriptomic analysis of allergen-specific T cells in allergy and asthma. Sci. Immunol. 5 (2020), doi:10.1126/SCIIMMUNOL.ABA6087.

22. L. Qiu, R. Lai, Q. Lin, E. Lau, D. M. Thomazy, D. Calame, R. J. Ford, L. W. Kwak, R. A. Kirken, H. M. Amin, Autocrine release of interleukin-9 promotes Jak3-dependent survival of ALK+ anaplastic large-cell lymphoma cells. Blood. 108, 2407–2415 (2006).

23. M. Fischer, M. Bijman, D. Molin, F. Cormont, C. Uyttenhove, J. van Snick, C. Sundström, G. Enblad, G. Nilsson, Increased serum levels of interleukin-9 correlate to negative prognostic factors in Hodgkin’s lymphoma. Leukemia. 17, 2513–2516 (2003).

24. R. Purwar, C. Schlapbach, S. Xiao, H. S. Kang, W. Elyaman, X. Jiang, A. M. Jetten, S. J. Khoury, R. C. Fuhlbrigge, V. K. Kuchroo, R. A. Clark, T. S. Kupper, Robust tumor immunity to melanoma mediated by interleukin-9-producing T cells. Nat. Med. 18, 1248–53 (2012).

25. Y. Lu, S. Hong, H. Li, J. Park, B. Hong, L. Wang, Y. Zheng, Z. Liu, J. Xu, J. He, J. Yang, J. Qian, Q. Yi, Th9 cells promote antitumor immune responses in vivo. J. Clin. Invest. 122, 4160–71 (2012).

26. S. Rauber, M. Luber, S. Weber, L. Maul, A. Soare, T. Wohlfahrt, N. Y. Lin, K. DIetel, A. Bozec, M. Herrmann, M. H. Kaplan, B. Weigmann, M. M. Zaiss, U. Fearon, D. J. Veale, J. D. Cañete, O. DIstler, F. Rivellese, C. Pitzalis, M. F. Neurath, A. N. J. McKenzie, S. Wirtz, G. Schett, J. H. W. DIstler, A. Ramming, Resolution of inflammation by interleukin-9-producing type 2 innate lymphoid cells. Nat. Med. 23, 938–944 (2017).

27. P. A. Reche, The tertiary structure of γc cytokines dictates receptor sharing. Cytokine. 116, 161–168 (2019).

28. S. Singh, N. Chaturvedi, G. Rai, De novo modeling and structural characterization of IL9-IL9 receptor complex: a potential drug target for hematopoietic stem cell therapy. Netw. Model. Anal. Heal. Informatics Bioinforma. 9, 1–10 (2020).

29. K. Verstraete, S. Koch, S. Ertugrul, I. Vandenberghe, M. Aerts, G. Vandriessche, C. Thiede, S. N. Savvides, Efficient production of bioactive recombinant human Flt3 ligand in E. coli. Protein J. 28, 57–65 (2009).

30. T. Evangelidis, S. Nerli, J. Nováček, A. E. Brereton, P. A. Karplus, R. R. Dotas, V. Venditti, N. G. Sgourakis, K. Tripsianes, Automated NMR resonance assignments and structure determination using a minimal set of 4D spectra. Nat. Commun. 9, 1–13 (2018).

31. J. L. Boulay, W. E. Paul, Hematopoietin sub-family classification based on size, gene organization and sequence homology. Curr. Biol. 3, 573–581 (1993).

32. K. Verstraete, L. van Schie, L. Vyncke, Y. Bloch, J. Tavernier, E. Pauwels, F. Peelman, S. N. Savvides, Structural basis of the proinflammatory signaling complex mediated by TSLP. Nat. Struct. Mol. Biol. 21, 375–82 (2014).

33. K. Verstraete, F. Peelman, H. Braun, J. Lopez, D. Van Rompaey, A. Dansercoer, I. Vandenberghe, K. Pauwels, J. Tavernier, B. N. Lambrecht, H. Hammad, H. De Winter, R. Beyaert, G. Lippens, S. N. Savvides, Structure and antagonism of the receptor complex mediated by human TSLP in allergy and asthma. Nat. Commun. 8, 1–17 (2017).

34. S. R. Sprang, J. Fernando Bazan, Cytokine structural taxonomy and mechanisms of receptor engagement. Current opinion in structural biology 1993, 3:815-827. Curr. Opin. Struct. Biol. 3, 815–827 (1993).

35. D. A. Rozwarski, A. M. Gronenborn, G. M. Clore, J. F. Bazan, A. Bohm, A. Wlodawer, M. Hatada, P. A. Karplus, Structural comparisons among the short-chain helical cytokines. Structure. 2, 159–173 (1994).

36. H. W. Park, K. G. Tantisira, Genetic signatures of asthma exacerbation. Allergy, Asthma Immunol. Res. 9, 191–199 (2017).

37. S. Moretti, G. Renga, V. Oikonomou, C. Galosi, M. Pariano, R. G. Iannitti, M. Borghi, M. Puccetti, M. De Zuani, C. E. Pucillo, G. Paolicelli, T. Zelante, J.-C. Renauld, O. Bereshchenko, P. Sportoletti, V. Lucidi, M. C. Russo, C. Colombo, E. Fiscarelli, C. Lass-Flörl, F. Majo, G. Ricciotti, H. Ellemunter, L. Ratclif, V. N. Talesa, V. Napolioni, L. Romani, A mast cell-ILC2-Th9 pathway promotes lung inflammation in cystic fibrosis. Nat. Commun. 8, 1–13 (2017).

38. H. Aschard, E. Bouzigon, E. Corda, A. Ulgen, M. H. Dizier, F. Gormand, M. Lathrop, F. Kauffmann, F. Demenais, Sex-specific effect of IL9 polymorphisms on lung function and polysensitization. Genes Immun. 10, 559–565 (2009).

39. A. Schuurhof, L. Bont, C. L. E. Siezen, H. Hodemaekers, H. C. Van Houwelingen, T. G. Kimman, B. Hoebee, J. L. L. Kimpen, R. Janssen, Interleukin-9 polymorphism in infants with respiratory syncytial virus infection: An opposite effect in boys and girls. Pediatr. Pulmonol. 45, 608–613 (2010).

40. A. Pasvenskaite, R. Liutkeviciene, G. Gedvilaite, A. Vilkeviciute, V. Liutkevicius, V. Uloza, The Role of IL-9 Polymorphisms and Serum IL-9 Levels in Carcinogenesis and Survival Rate for Laryngeal Squamous Cell Carcinoma. Cells. 10 (2021), doi:10.3390/cells10030601.

41. Y. Kimura, T. Takeshita, M. Kondo, N. Ishii, M. Nakamura, J. Van Snick, K. Sugamura, Sharing of the IL-2 receptor gamma chain with the functional IL-9 receptor complex. Int. Immunol. 7, 115–20 (1995).

42. P. Gonnord, B. R. Angermann, K. Sadtler, E. Gombos, P. Chappert, M. Meier-Schellersheim, R. Varma, A hierarchy of affinities between cytokine receptors and the common gamma chain leads to pathway cross-talk. Sci. Signal. 11 (2018), doi:10.1126/scisignal.aal1253.

43. A. T. Waickman, H. R. Keller, T. H. Kim, M. A. Luckey, X. Tai, C. Hong, C. Molina-París, S. T. R. Walsh, J. H. Park, The Cytokine Receptor IL-7Rα Impairs IL-2 Receptor Signaling and Constrains the In Vitro Differentiation of Foxp3+ Treg Cells. iScience. 23 (2020), doi:10.1016/j.isci.2020.101421.

44. A. R. Aricescu, W. Lu, E. Y. Jones, A time- and cost-efficient system for high-level protein production in mammalian cells. Acta Crystallogr. Sect. D Biol. Crystallogr. 62, 1243–1250 (2006).

45. C. Basilico, A. Hultberg, C. Blanchetot, N. de Jonge, E. Festjens, V. Hanssens, S.-I. Osepa, G. De Boeck, A. Mira, M. Cazzanti, V. Morello, T. Dreier, M. Saunders, H. de Haard, P. Michieli, Four individually druggable MET hotspots mediate HGF-driven tumor progression. J. Clin. Invest. 124, 3172 (2014).

46. de H. HJ, van N. N, R. A, H. SE, R. RC, H. P, de B. AP, A. JW, H. HR, A large non-immunized human Fab fragment phage library that permits rapid isolation and kinetic analysis of high affinity antibodies. J. Biol. Chem. 274, 18218–18230 (1999).

47. P. Güntert, Automated structure determination from NMR spectra. Eur. Biophys. J. 38 (2009), pp. 129– 143.

48. Y. Shen, F. Delaglio, G. Cornilescu, A. Bax, TALOS+: A hybrid method for predicting protein backbone torsion angles from NMR chemical shifts. J. Biomol. NMR. 44, 213–223 (2009).

49. J. P. Linge, M. A. Williams, C. A. E. M. Spronk, A. M. J. J. Bonvin, M. Nilges, Refinement of protein structures in explicit solvent. Proteins Struct. Funct. Bioinforma. 50, 496–506 (2003).

50. A. J. Nederveen, J. F. Doreleijers, W. Vranken, Z. Miller, C. A. E. M. Spronk, S. B. Nabuurs, P. Güntert, M. Livny, J. L. Markley, M. Nilges, E. L. Ulrich, R. Kaptein, A. M. J. J. Bonvin, RECOORD: A recalculated coordinate database of 500+ proteins from the PDB using restraints from the BioMagResBank. Proteins Struct. Funct. Bioinforma. 59, 662–672 (2005).

51. A. Bhattacharya, R. Tejero, G. T. Montelione, Evaluating protein structures determined by structural genomics consortia. Proteins Struct. Funct. Bioinforma. 66, 778–795 (2006).

52. W. Kabsch, XDS. Acta Crystallogr. D. Biol. Crystallogr. 66, 125–32 (2010).

53. A. J. McCoy, R. W. Grosse-Kunstleve, P. D. Adams, M. D. Winn, L. C. Storoni, R. J. Read, Phaser crystallographic software. J. Appl. Crystallogr. 40, 658–674 (2007).

54. P. Emsley, B. Lohkamp, W. G. Scott, K. Cowtan, Features and development of Coot. Acta Crystallogr. Sect. D Biol. Crystallogr. 66, 486–501 (2010).

55. P. D. Adams, P. V. Afonine, G. Bunkóczi, V. B. Chen, I. W. Davis, N. Echols, J. J. Headd, L. W. Hung, G. J. Kapral, R. W. Grosse-Kunstleve, A. J. McCoy, N. W. Moriarty, R. Oeffner, R. J. Read, D. C. Richardson, J. S. Richardson, T. C. Terwilliger, P. H. Zwart, PHENIX: A comprehensive Python-based system for macromolecular structure solution. Acta Crystallogr. Sect. D Biol. Crystallogr. 66, 213–221 (2010).

56. P. W. Bricogne G., Blanc E., Brandl M., Flensburg C., Keller P., W. T. O. Roversi P, Sharff A., Smart O.S., Vonrhein C., BUSTER version 2.10.3 (2017).

57. R. P. Joosten, F. Long, G. N. Murshudov, A. Perrakis, The PDB-REDO server for macromolecular structure model optimization. IUCrJ. 1, 213–220 (2014).

58. Y. Li, R. Sousa, Expression and purification of E. coli BirA biotin ligase for in vitro biotinylation. Protein Expr. Purif. 82, 162–167 (2012).

59. X. Robert, P. Gouet, Deciphering key features in protein structures with the new ENDscript server. Nucleic Acids Res. 42, 320–324 (2014).

